# Chemo-mechanical Diffusion Waves Orchestrate Collective Dynamics of Immune Cell Podosomes

**DOI:** 10.1101/2021.11.23.469591

**Authors:** Ze Gong, Koen van den Dries, Alessandra Cambi, Vivek B. Shenoy

## Abstract

Immune cells, such as macrophages and dendritic cells, can utilize podosomes, actin-rich protrusions, to generate forces, migrate, and patrol for foreign antigens. In these cells, individual podosomes exhibit periodic protrusion and retraction cycles (vertical oscillations) to probe their microenvironment, while multiple podosomes arranged in clusters demonstrate coordinated wave-like spatiotemporal dynamics. However, the mechanisms governing both the individual vertical oscillations and the coordinated oscillation waves in clusters remain unclear. By integrating actin polymerization, myosin contractility, actin diffusion, and mechanosensitive signaling, we develop a chemo-mechanical model for both the oscillatory growth of individual podosomes and wave-like dynamics in clusters. Our model reveals that podosomes show oscillatory growth when the actin polymerization-associated protrusion and the signaling-associated myosin contraction occur at similar rates, while the diffusion of actin monomers within the cluster drives mesoscale coordination of individual podosome oscillations in an apparent wave-like fashion. Our model predicts the influence of different pharmacological treatments targeting myosin activity, actin polymerization, and mechanosensitive pathways, as well as the impact of the microenvironment stiffness on the wavelengths, frequencies, and speeds of the chemo-mechanical waves. Overall, our integrated theoretical and experimental approach reveals how collective wave dynamics arise due to the coupling between chemo-mechanical signaling and actin diffusion, shedding light on the role of podosomes in immune cell mechanosensing within the context of wound healing and cancer immunotherapy.

## Introduction

Immune cells, such as dendritic cells (DCs) and macrophages, can crawl between cells and components of the extracellular matrix (ECM) to patrol for foreign antigens. During this process, these cells utilize actin-rich protrusive structures called podosomes to control adhesions, degrade the surrounding ECM, and remodel the extracellular environment^1-3^. Podosomes are characterized by an actin-based core surrounded by an adhesive ring consisting of integrins and adaptor proteins, such as vinculin and talin^4,5^. Podosomes generate protrusive forces to penetrate into the underlying ECMs at the core, while applying tensile forces to pull the matrices at the rings^6,7^. Studies in DCs and macrophages have shown that individual podosomes exhibit periodic oscillation in the fluorescence intensity of core actin, ring components, and the protrusive forces exerted at the podosome core^7-9^. Collectively, podosomes in immune cells are organized in clusters and show spatially correlated behaviors, where the oscillations of podosome components travel in a wave-like manner within the cluster^10,11^. However, it remains unclear how the core protrusive forces and ring tensile forces lead to the oscillations of individual podosomes and how these vertical oscillations are coordinated to form spatiotemporal wave patterns in clusters. Understanding these processes can provide insights on how podosomes in a cluster collectively probe and respond to chemo-mechanical cues from their surroundings, which is essential for their function.

To obtain biophysical insights into the dynamics of podosome clusters, we adopted a bottom-up modelling approach. We first integrated actin polymerization, myosin contractility, and mechano-sensitive signaling pathways into a chemo-mechanical model for oscillatory growth of individual podosomes. By considering diffusion of actin monomers within a cluster, our model demonstrates how correlations can arise between the vertical oscillations of podosomes to form coordinated wave dynamics in clusters, which we call chemo-mechanical diffusion waves. Next, using fluorescence microscopy and spatiotemporal image correlation spectroscopy in living primary human DCs, we extracted the wavelengths, oscillation periods, and speeds of podosome wave dynamics. Our model is quantitatively validated by predicting the influence of pharmacological treatments targeting force-generating processes (myosin activity, actin polymerization, Rho and Rho-associated kinase (Rho-ROCK) pathways) as well as the impact of substrate stiffness on the podosome dynamics. The mechanistic understanding of how the individual podosome oscillations, driven by chemo-mechanical signaling, are coordinated at the mesoscale and lead to wave-like propagation of podosome components and forces, provides means to modulate immune cell mechanosensing and migration within the context of wound healing and cancer immunotherapy.

## Results

### A chemo-mechanical model for dynamics of individual podosomes

To develop a biophysical description of podosome growth, we start by considering the molecular mechanisms of force generation in a podosome. Podosomes have a conical structure characterized by a protrusive actin-rich core, an adhesive integrin ring, and ventral actin filaments that connect the core with the ring^12,13^ (Fig. 1A-B). During podosome growth, actin monomers (G-actin) continuously polymerize into actin filaments (F-actin), generating a protrusive (or compressive) force in the core to drive core growth. At the same time, myosin motors are dynamically recruited to the ventral actin filaments, generating an active contractile force to constrain growth^3,9,12^. The core protrusive force generated by actin polymerization *F*_*p*_ is balanced by the tensile force generated by the actomyosin contractility *F*_*r*_ according to *F*_*p*_=*F*_*r*_ cos(θ), where we assume θ to be a constant angle between the ventral actin filaments and the core F-actin (Fig.1A). This assumption yields a geometric constraint between the ventral actin filament length *x*and the core height above the undeformed substrate *l*, i.e., *xcos*(θ) =*l*. Next, we describe the dynamics of the two force-generating processes individually:

**Fig 1.**
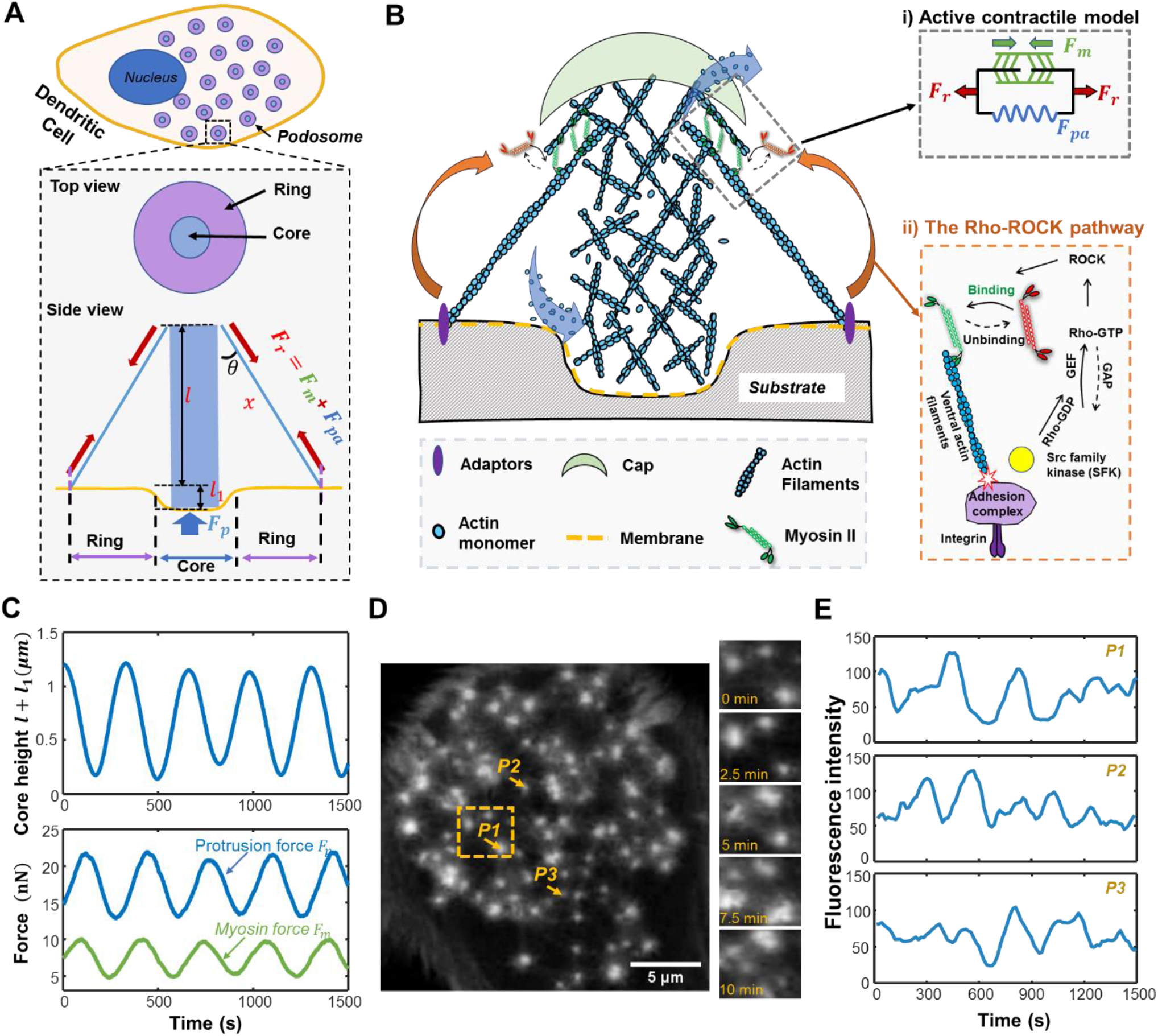
A chemo-mechanical model for the oscillatory growth of individual podosomes. (A) Schematic showing a dendritic cell with podosomes. We also show simplified geometry of the podosome with top and side views. (B) Schematic describing the chemo-mechanical model for podosome growth. The orange arrows indicate mechano-sensitive signaling through the Rho-ROCK pathway. Inset i): the active contractile model for myosin-based force generation, where the active contractile forces *F*_*m*_ (the green element with arrows), the passive actin filament elastic force *F*_*pa*_ (blue spring), and total ring force *F*_*r*_ (red arrows) are marked. Inset ii): tensile forces in the ring trigger a conformation change in vinculin, exposing binding sites of Src family of tyrosine kinases (SFKs); this change promotes Rho-GTPases by controlling the activity of guanine nucleotide exchange factors (GEFs) and GTPase-activating proteins (GAPs), which eventually increases the level of contractile forces in the ventral F-actins (C) Representative curves showing the theoretically predicted dynamics for podosome core height *l* + *l*_1_ (top panel), protrusion force *F*_*p*_(blue line in the bottom panel), and myosin force *F*_*m*_ (green line in the bottom panel). (D) A representative Lifeact-RFP transfected dendritic cell. Podosome dynamics at different times for the yellow-dashed region are shown in the top panel. Scale bar 5 μm. (E) Dynamics of Lifeact-RFP fluorescence intensity for three representative podosomes, which are marked in (D).

#### Actin polymerization drives the protrusion of podosome cores

In the podosome core, the protrusive force *F*_*p*_ generated by actin polymerization deforms the underlying substrate following *F*_*p*_=*k*_*s*_*l*_1_, where *k*_*s*_ and *l*_1_ are the stiffness and the displacement of the substrate, respectively. The resistance force from the substrate reduces actin polymerization^14,15^, which can be expressed as *V*_*p*_=*V*_*pm*_(1 −*F*_*p*_/*F*_*sp*0_), where *V*_*pm*_ is the maximum polymerization speed and *F*_*sp*0_ is the characteristic protrusive force generated by the core. Note that, to account for larger protrusive forces on stiffer substrates revealed by protrusion force microcopy^7^, we write the characteristic stall force as *F*_*sp*0_ =*F*_*p*0_*k*_*s*_/(*k*_*c*_ + *k*_*s*_), where *k*_*c*_ is the core stiffness and *F*_*p*0_ is the characteristic protrusive force on rigid substrate; the protrusive force increases linearly with the substrate stiffness at low levels but saturates on very stiff substrates. The growth rate of the core height is determined by the difference between the polymerization *V*_*p*_ and depolymerization speed *V*_*d*_, which can be written as:

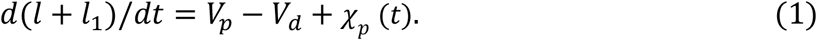

Here, *l* + *l*_1_ is the total podosome height, where *l* is the core height above the undeformed substrate, and the Gaussian noise *χ*_*p*_(*t*) accounts for fluctuations in the polymerization process (Fig.1A).

#### Mechano-sensitive recruitment of myosin to the ventral actin filaments

As the F-actin network assembles in the podosome core and generates a protrusive force, myosin motors exert contractile forces on the ventral actin filaments to balance the core protrusive force and constrain core growth. To model the contraction of actomyosin filaments, we adopt a two-element active contraction model^16^, consisting of an active element (with contractile force *F*_*m*_) in parallel with a passive elastic element (with stiffness *k*_*f*_). The active element characterizes myosin contractility, while the passive (elastic) element represents the stiffness of the ventral actin filaments (Fig. 1B, inset i). Thus, the tensile force sustained by the ring *F*_*r*_ can be written as the sum of the active and passive forces: *F*_*r*_=*F*_*m*_+ *k*_*f*_(*x*−*x*_0_),where *x*and *x*_0_ denote the current and initial length of the ventral actin filaments, respectively. In addition, myosin activity is known to be mediated by the Rho-ROCK pathway^10,17,18^. Specifically, the tensile force *F*_*r*_ transmitted to the substrate through the integrin adhesions can positively feed back to the active force *F*_*m*_ through this pathway^16^ (Fig. 1A, inset ii). In analogy with muscle fibers, the contraction of the actomyosin filaments, *x*−*x*_0_, reduces available binding sites for myosin, causing a decrease of bound myosin motors and active contractility^16,19,20^. Here, we use *α*to characterize the effects of feedback (proportional to the tensile force *F*_*r*_) and *γ*to characterize the effects of the actomyosin filament length on active contractility (refer to SI Note 1), which leads to the following equation governing the dynamics of myosin force:

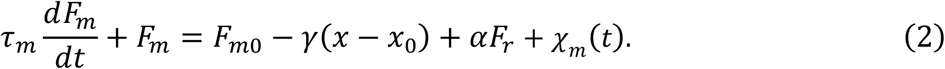

Here *τ*_*m*_ is the characteristic time for the myosin turnover, *F*_*m*0_ is the base level myosin force, and *χ*_*m*_(*t*) is a Gaussian noise signal accounting for fluctuations in myosin dynamics. The equations along with the parameters used in the simulations are summarized in the *Models and Methods* and Table S1.

### Actin polymerization and myosin recruitment govern podosome oscillatory growth

By combining the governing equations for the core protrusion and ring contraction dynamics (Eqs. 1&2), we can simulate the oscillatory behaviors of the core height, the protrusive force, and active myosin contractility (Fig. 1C). The time-averaged magnitudes of the simulated protrusive force (∼10 *nN*), ventral actin filament length (∼1 *μm*), and core height (∼700 *nm*) agree with previous experimental measurements^7,9,12^. To understand the mechanisms governing podosome oscillations, we identify four stages of a typical protrusion-retraction oscillatory cycle (Fig. 2A-B):

**Fig 2.**
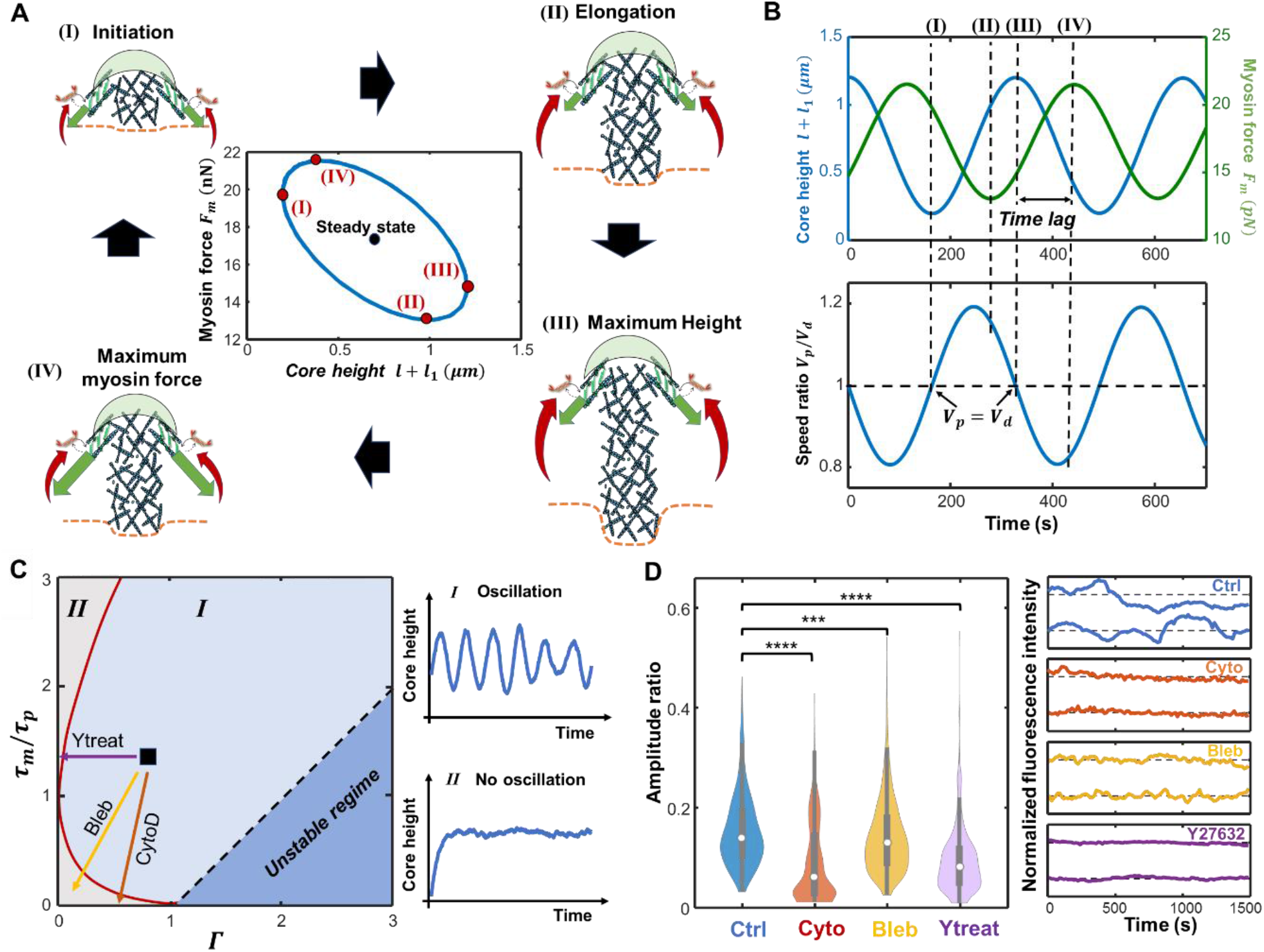
Model predicts the effects of pharmacological treatments on podosome oscillations. (A) Schematic showing the interplay of intracellular processes in the four stages of oscillatory podosome growth. The substrate provides only a small resistance force at the beginning, and actin monomers polymerize to drive podosome growth (stage I). As the podosome grows and deforms the substrate, the protrusion speed gradually decreases (stage II). The active myosin contractility (green arrows) continues to increase when the podosome core reaches its maximum height (stage III) because of the time delay associated with signaling feedback (red arrows). The podosome begins to retract in response to high levels of contractile forces (stage IV). The inset (middle panel): myosin force versus core height. (B) The core length *l* + *l*_1_ (blue line in top panel), myosin contractile force *F*_*m*_(green line in top panel), and the ratio between polymerization and depolymerization speed *V*_*p*_/*V*_*d*_ (blue line in bottom panel) plotted as a function of time. (C) Phase diagram showing two types of protrusion patterns: oscillatory protrusions (regime I) and monotonically growing protrusions (regime II), based on the ratio *τ*_*m*_/*τ*_*p*_ and the feedback parameter *Γ*. The arrows indicate the influence of cytochalasin D (CytoD, red), blebbistatin (Bleb, yellow), and Y27632 (Ytreat, purple) treatments on the dynamics. (D) (Left panel) The amplitude ratio, *r*_*a*_ for control (Ctrl, blue), cytochalasin D (red), Blebbistatin (yellow), and Y27632 (purple) treatments. Statistically significant differences are indicated (***p < 0.001, ****p < 0.0001, ANOVA with Benjamini-Hochberg procedure). (Right panel) Representative fluorescence intensity profiles for control and different pharmacological treatments.

I. In the early stage of growth, the podosome has a small core height and low levels of active myosin contractility (green arrows in Fig.2A). The podosome height increases as the polymerization speed exceeds the depolymerization speed (via Eq. 1).
II. As the F-actin network within the core grows and pushes against the substrate, the protrusive force *F*_*p*_ increases, resisting actin polymerization. The protrusion speed gradually decreases.
III. When the protrusion speed decreases to zero, the podosome height, protrusive force *F*_*p*_, and ring adhesion force *F*_*r*_ attain their maximum values, leading to a high level of feedback via the Rho-Rock pathway (red arrows in Fig.2A). In response to this signaling response, more myosin motors are recruited, generating larger myosin contractile forces. Since the myosin force requires the characteristic time *τ*_*m*_ to accumulate (via Eq. 2), there is a time delay between the maximum podosome height and the maximum myosin force.
IV. As the protrusion speed continues to decrease and drops below zero, the podosome starts retracting. Subsequently, the myosin force decays from its maximum value, and polymerization speed increases. Once the polymerization speed exceeds the depolymerization speed, the next oscillation cycle begins.

The two force-generating processes, i.e., the polymerization-associated protrusion and the signaling-associated myosin contraction, govern the dynamics of podosome oscillations (Fig. 2A). Ring contractile forces continue to increase when podosomes reach their maximum height, and podosomes begin to retract in response to high contractile forces in the ring. When contractile forces decrease and polymerization dominates, podosomes start to grow again. This back-and-forth interaction between the two force-generating processes regulates podosome oscillations. By transfecting human DCs with Lifeact-RFP and vinculin-GFP, we found that both the F-actin and vinculin fluorescence intensity oscillate periodically with time (Fig. 1D-E and Movie S1). It is worth noting that fluorescence traces of F-actin and vinculin are highly correlated (Fig. S1A-B). In the next section, we study the timescales governing the two competing force generation processes and how they affect the podosome dynamics.

### Phase diagram predicts the effects of pharmacological treatments on podosome dynamics

We applied a linear perturbation analysis to study the stability of the steady-state solutions for the core protrusion and the actomyosin contractility of the podosomes (refer to *Models and Methods* section). We found that podosome oscillations are governed by the signaling-associated feedback parameter 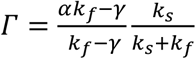 and the ratio *τ*_*m*_ /*τ*_*p*_ between the timescales governing myosin turnover (*τ*_*m*_) and the core protrusion (*τ*_*p*_). Note here the protrusion timescale is 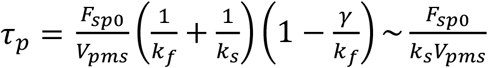, which characterizes the time it takes a podosome with polymerization speed *V*_*pms*_ to grow by a characteristic substrate displacement *F*_*sp*0_/*k*_*s*_. The magnitude of the feedback parameter *Γ*∼ *α*,characterizes the strength of Rho-ROCK pathway when the stiffness for the passive component is much larger than the effective stiffness of the active element of actomyosin filaments, i.e., *k*_*f*_ ≫ *α*. This stability analysis allows us to generate a phase diagram, which reveals distinct phases of oscillatory or monotonic dynamic behavior based on the feedback parameter *Γ* and the timescale ratio *τ*_*m*_/*τ*_*p*_(Fig. 2C). We find that podosomes exhibit oscillatory growth (region I in Fig. 2C) when the two timescales are comparable (*τ*_*m*_/*τ*_*p*_∼1,) and the feedback parameter *Γ* is at an intermediate value. When the two timescales substantially differ, the system relaxes monotonically to the steady-state podosome length (region II in Fig. 2C); this indicates that oscillatory growth requires that the two competing force-generating mechanisms occur at similar rates. At the same time, the mechanosensitive Rho-ROCK pathway is critical for oscillations, since a very weak signaling feedback *Γ* → 0 also leads to monotonic podosome growth. Linear stability analysis gives the oscillation period 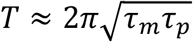, which is on the order of several minutes in line with our experiments (Fig. 1E) and previous findings^9,10^.

We can further use the phase diagram to predict the effects of pharmacological treatments on podosome dynamics. By using different treatments targeting the force generation processes (i.e., actin polymerization and myosin contractility) or mechanosensitive Rho-ROCK pathways, our model predicts that podosomes exhibit non-oscillatory behaviors (arrows in Fig. 2C and S2). This is because the two timescales (*τ*_*m*_,*τ*_*p*_) become not comparable or signaling feedback *Γ* is reduced (for details refer to SI Note 2). To test these predictions, we treated DCs with cytochalasin D, blebbistatin, and Y26732 to inhibit actin polymerization, myosin contractility, and the Rho-ROCK signaling, respectively. By evaluating the oscillation amplitude ratio of the Lifeact-RFP fluorescence intensity (i.e., the ratio between the oscillation amplitude and the time-averaged intensity^21^) to characterize the relative oscillation intensity, we found that all these treatments significantly reduce the amplitude ratio. This reduced ratio indicates that oscillations are inhibited after pharmacological treatments, in agreement with our predictions (Fig. 2C-D). Overall, the combined experimental and theoretical analysis demonstrate that podosomes oscillate only when the polymerization-associated protrusion and signaling-associated myosin contraction occur at similar rates.

### Actin diffusion drives coordinated wave-like patterns in podosome clusters

Next, we studied how the oscillations of individual podosomes give rise to wave-like patterns in the collective dynamics of podosomes. To model these spatiotemporal collective dynamics, we consider the diffusion and exchange of actin within the cluster. The time evolution of G-actin concentration *c*_*a*_(*x,y,t*) in the plane of the substrate at time *t* is determined by both actin diffusion and actin polymerization (or depolymerization), which is written as:

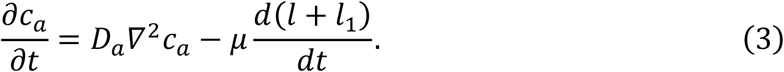

Here *D*_*a*_ is the G-actin diffusion constant and parameter *μ* controls the drop of G-actin concentration as actin monomers are assembled into the core to increase its height *l* + *l*_1_; this is proportional to the cross-section area *A* and the planar density of actin monomers *ρ* of podosome core, i.e., *μ* ∝ *Aρ*. At the same time, a larger G-actin concentration at the podosome core increases the polymerization speed, that is *V*_*pm*_=*V*_*p*0_ + *βc*_*a*_, where a linear dependence of polymerization speed on actin concentration with sensitivity *β* is assumed for simplicity. By integrating this diffusion process with the model for individual podosome dynamics (see *Models and Methods* section), we can simulate the spatiotemporal dynamics of the podosome cluster using discrete and coarse-grained approaches. We first consider the dynamics of an array of podosomes individually with a uniform distance *d*_0_ from each other in the discrete model (Fig. 3A and S3A); we then further generalize the discrete model using a coarse-grained (or continuum) approach (Fig. 3B and S3B), where only a spatial average of the podosome heights, treated as a continuous variable, is studied (refer to *Models and Methods* section and *SI Note 2*). By assuming an initial distribution where only podosomes at the center have large heights, both the discrete and continuum models show that a height wave front originates from the center and gradually transits outward, forming a radial wave pattern (Movies S2-S3). These simulated radial patterns closely resemble the observed LifeAct-RFP intensity dynamics in the experiments (Fig. 3C).

**Fig 3.**
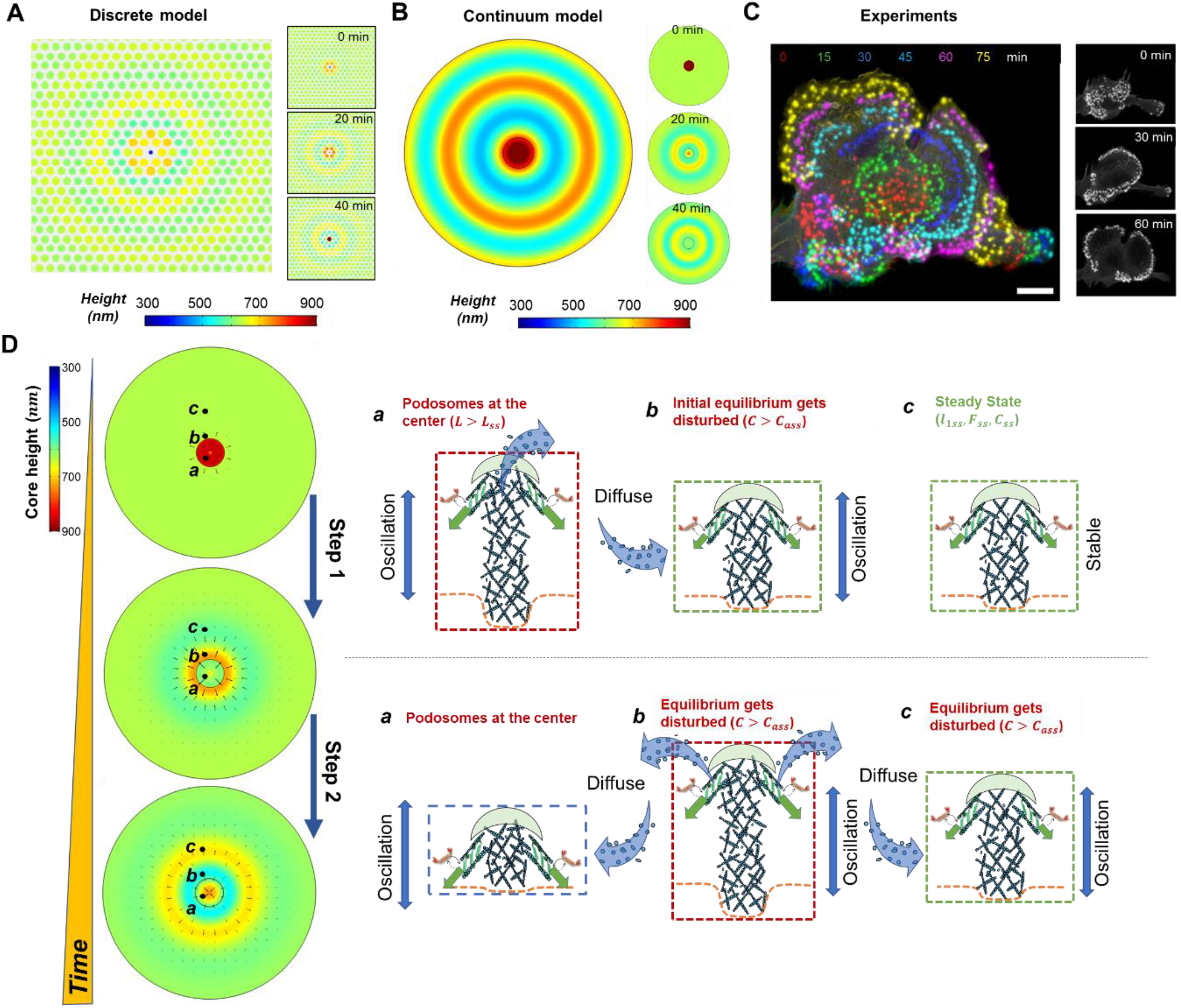
Mechanism of radial wave formation in podosome clusters. (A-B) The simulated heights in the podosome cluster using (A) a discrete model of podosomes and (B) a continuum model showing radial wave patterns. Each circle in panel (A) represents an individual podosome with the size and color representing its height. Different colors in panel (B) represent the podosome heights. (C) DCs transfected with LifeAct-RFP showing the propagation of radial waves. Colors denote the fluorescence-labeled podosomes at different times (Figure adapted from our previous study ^10^). The insets in (A-C) show the radial wave patterns at different times. Scale bar 10 μm. (D) The schematic showing the mechanism for the propagation of radial waves. The black arrows in left panels indicate the flow of G-actin.

To understand the mechanism for the formation and propagation of radial waves, we take a closer look at the dynamics of individual podosomes located at different distances from the center during the wave propagation (Fig. 3D).

Phase 1: The podosomes at the center with large heights (above the steady-state core height, denoted with *a*) depolymerize, release G-actin, and increase the local G-actin concentration. Then, G-actin diffuses outward, altering the G-actin concentration near podosome *b* located away from the center, thus disrupting the previous balance between actin consumption and release. The podosome *b* starts growing in turn.

Phase 2: As the height of podosome *a* reduces and reaches its minimum value, podosome *b* reaches its maximum height and begins to depolymerize, sending G-actin further outwards to activate the oscillation of podosome *c*. The radial wave pattern forms and propagate.

Overall, the oscillations of individual podosomes constantly change the local actin concentration, and the uneven spatial distribution of actin concentration drives diffusion in the podosome cluster, leading to the wave patterns in podosome core heights. As this phenomenon involves coordination of actin diffusion, mechanical forces, and biochemical signaling, we have coined the term *chemo-mechanical diffusion waves* to describe the collective dynamics of podosome clusters.

### Theory quantitatively predicts the periods and wavelengths of chemo-mechanical waves

Most chemo-mechanical diffusion waves observed in experiments show some degree of randomness in the wave patterns. To simulate the random waves (Movie S1), we assume a random distributed podosome heights in the beginning. Both the discrete and continuum approaches capture the propagation of random waves in the core heights of podosome clusters (Fig. 4A, 4D, S3, and Movies S2-S3). Plots of the dynamics of podosome core heights from the discrete model show that the dynamics of neighboring podosomes are strongly correlated, while the dynamics of distant podosomes are generally uncorrelated (Fig. 4B). This prediction is further supported by our experiments based on LifeAct-RFP intensity traces of neighboring and distant podosome pairs in the cluster (Fig. 4C). Furthermore, by using the kymograph analysis (which displays a series of images of the indicated rectangular area in Fig.4D-E over time), we found that the simulated peaks of podosome heights travel within the cluster, forming random wave patterns (Fig. 4D). The simulated patterns span regions with a typical length scale of ∼3 um and persist with a lifetime of ∼5 min, in agreement with the Life-Act fluorescence patterns in our experiments (Fig. 4E). Similar to the radial waves, these random wave patterns originate from the individual podosome oscillations and actin diffusion. By applying a small perturbation (∼*e*^*iqr*+*iωt*^,refer to *Models and Methods*), we obtain the angular wavenumber of the chemo-mechanical diffusion waves:

**Fig 4.**
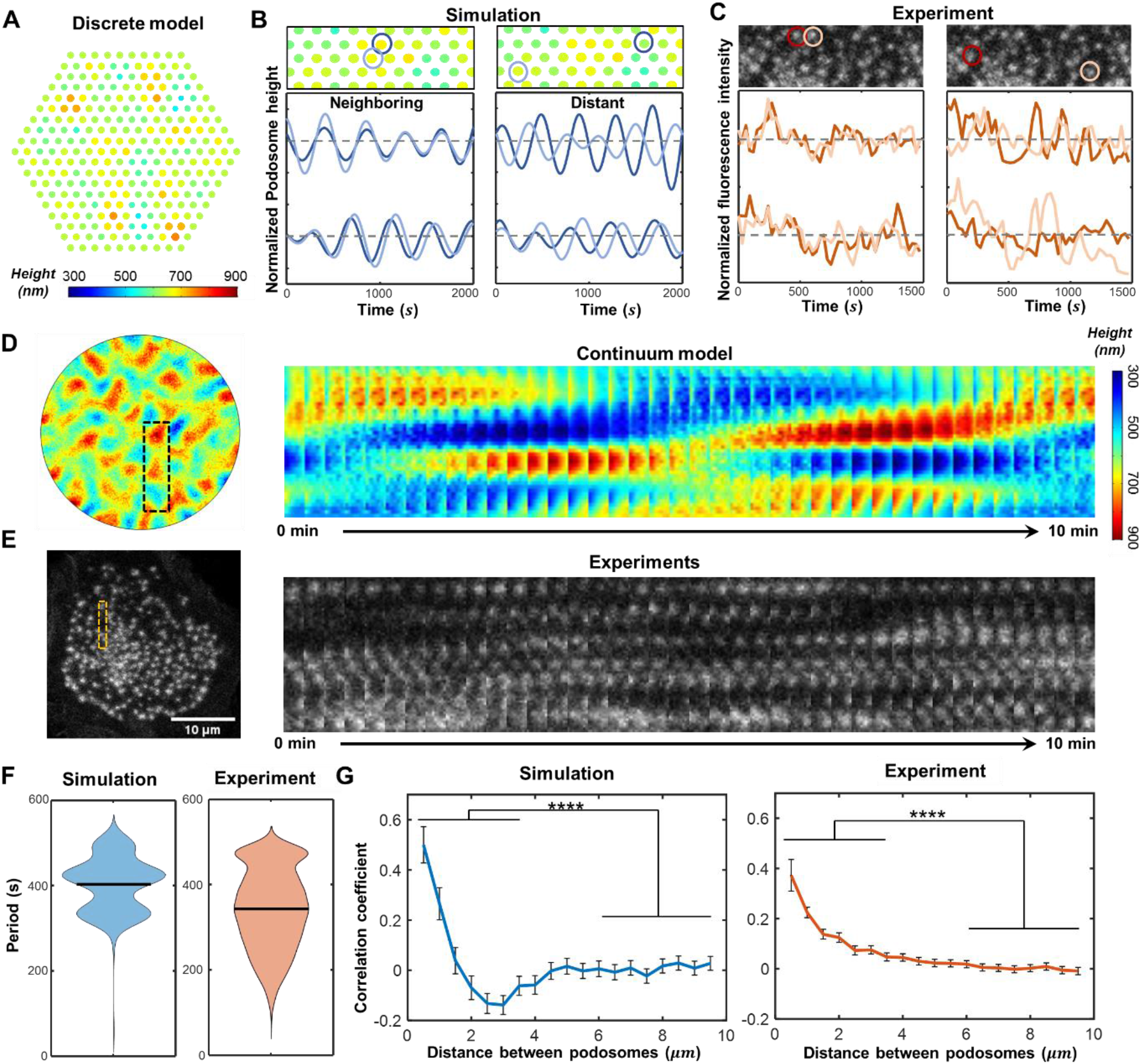
Model quantitively predicts the wavelengths and periods of random waves. (A) Simulated heights in the podosome cluster using the discrete approach showing random wave patterns. Each circle represents an individual podosome with its size and color representing height. (B-C) Plots of (B) simulated podosome heights and (C) LifeAct-RFP intensity over time for (left panel) two representative neighboring and (right panel) distant podosome pairs. The representative podosome pairs are marked in the movie snapshots in the top panels. Data are normalized by (B) mean height or (C) mean RFP intensity. (D) Simulated heights of a podosome cluster using the continuum approach with the kymograph of the indicated rectangular area. The color legend is the same as the legend in (A). (E) A representative LifeAct-RFP transfected DC with the kymograph of the indicated rectangular area. Scale bar 10 μm. (F) The oscillation periods of individual podosomes extracted by fast Fourier transformation for (left panel) simulations and (right panel) experimental results. (G) (Left panel) Average correlation coefficient of simulated core heights and (right panel) experimentally measured actin intensity fluctuations as a function of podosome pair distance (over 10^4^ podosome pairs of 11 DCs for experiments, over 10^4^ podosome pairs for the simulated podosome cluster). Error bars indicate standard deviation (****p < 0.0001, T-test).

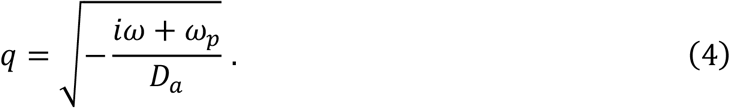

Here *ω* =*2π*/*T* is the radial frequency of podosome oscillations and *ω*_*p*_=*βμV*_*d*_/*V*_*pms*_ represents the effective exchange rate between the core F-actin and free G-actin. The real part of the wave number yields the wavelength *λ* =*2π*/*Re*(*q*), which can be approximated as a diffusion length scale 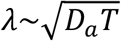 for a small exchange rate (*ω*_*p*_≪ *ω*). The imaginary part characterizes the damping effect within the system (refer to SI Note 5).

Next, to quantitatively compare our simulations with experiments, we extract the wave periods and wavelengths from our podosome simulations and LifeAct-RFP intensity dynamics. Using Fast Fourier Transformations (FFT) to process the experimentally measured LifeAct-RFP intensity fluctuations of individual podosomes, we extract the oscillation periods of all podosomes in a cluster (Fig. S4, refer to *Models and Methods*). We find that the average oscillation period extracted from experiments is between five and eight minutes (∼400 s), in line with our simulation results (Fig. 4F) and our analytically predicted period for individual podosome oscillation 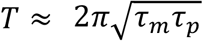. To extract the wavelength from the simulations and experiments, we calculate the correlation coefficient, which defines the correlation of intensity dynamics between two podosomes (refer to *Models and Methods*). We find the correlation coefficients extracted from both the experiments and the simulations decay with the distance between the podosome pair (Fig. 4G), indicating that podosome dynamics become gradually uncorrelated as the distance increases. From these correlation coefficient plots, we obtain the characteristic length scale for chemo-mechanical diffusion waves, which agrees with our model predicted wavelength scale *λ*∼3 *μm*.

### Oscillations of individual podosomes mediate the propagation of chemo-mechanical waves

Our model predicts that individual podosome oscillations cause spatial variation in G-actin density, which in turn drives chemo-mechanical diffusion waves. We next consider how wave propagation is impacted when oscillations of individual podosomes are inhibited. First, in our coarse-grained model, we disrupt the individual podosome oscillations by inhibiting myosin contractility or actin polymerization (as we have shown in Fig. 2C-D). We find that the inhibition of either contractility or polymerization disrupts the chemo-mechanical diffusion waves (Fig. 5A and S5A-B); this is because the G-actin concentration becomes spatially homogenous in steady state once individual podosome oscillations are inhibited and therefore cannot consume or release G-actin (Fig. 5A). To further quantify the changes of chemo-mechanical diffusion waves, we characterize the wave propagation speed using the phase velocity *v*_*c*_ = *λ*/*T* as a function of the oscillation period *T* and the effective actin exchange rate *ω*_*p*_(Fig. 5C, refer to *Models and Methods*). As a larger oscillation period *T* corresponds to less pronounced oscillations (*T* → ∞ means non-oscillatory growth), our model predicts that the wave propagation speed *v*_*c*_ decreases after the inhibition of individual podosome oscillations (arrows in Fig. 5C).

**Fig 5.**
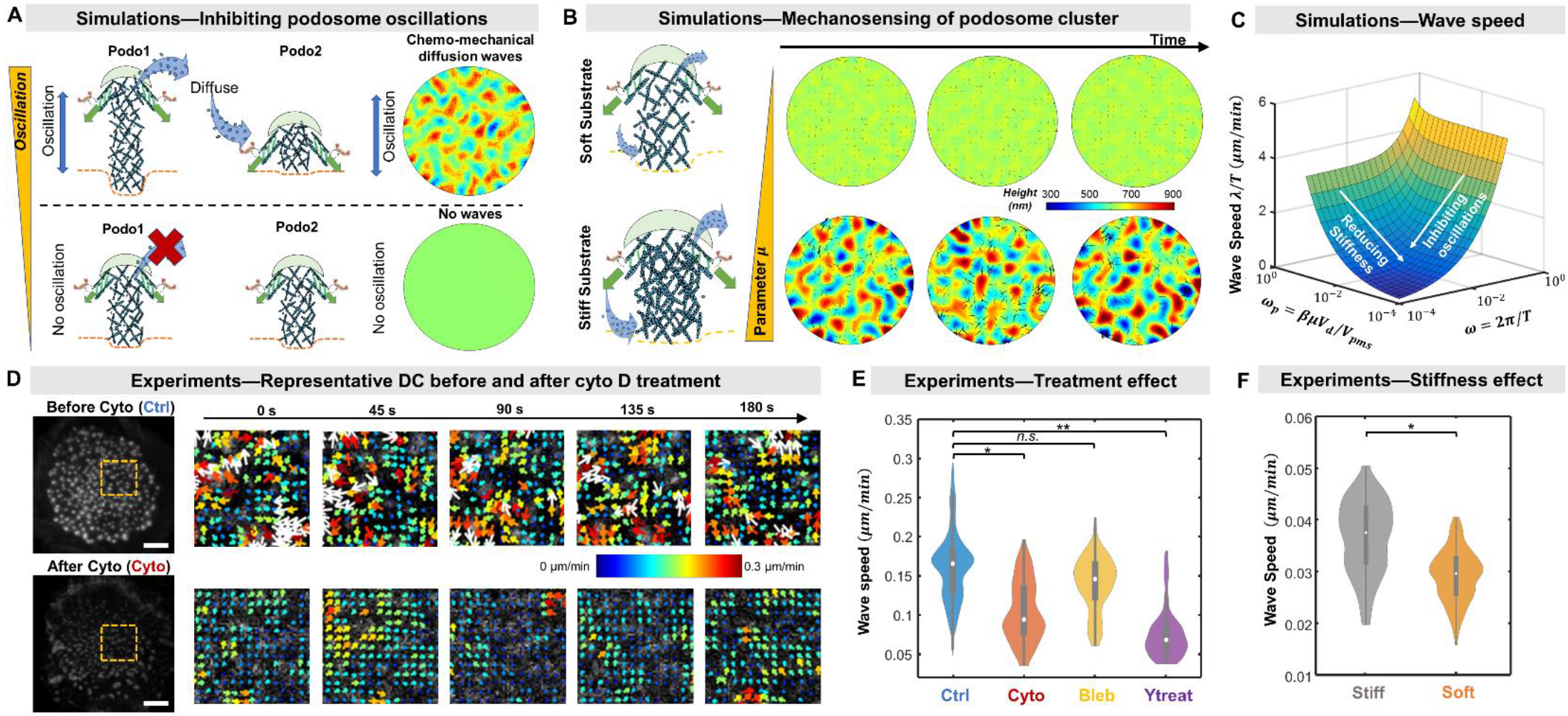
Model predicts the inhibition of pharmacological treatments and the influence of substrate stiffness on wave propagation. (A) (Left panels) Schematic and (right panels) simulations showing that inhibition of podosome oscillations disrupt chemo-mechanical diffusion waves. (B) The simulated podosome heights in a cluster plotted with time for (top panels) soft and (bottom panels) stiff substrates. The black arrows in the cluster indicate the magnitudes and directions of G-actin flow. (C) Wave propagation velocity *v*_*c*_ plotted versus the oscillation period *T* and diffusion constant *D*_*a*_. The arrow indicates the influence of pharmacological treatments that abrogate oscillations. (D) Representative DC (top panels) before and (right panels) after adding cytochalasin D. DCs are transfected with LifeAct-RFP using confocal microscopy with 15 s frame intervals. Time series subjected to twSTICS analysis are plotted as vector maps. The arrows indicate flow directions, and both the size and color denote flow magnitudes. Note that the arrows with white color indicate the magnitudes exceed 0.3 *μm*/*min*. (E) The velocity magnitudes for control (Ctrl, blue), cytochalasin D (Cyto, red), Blebbistatin (Bleb, yellow), and Y27632 (Ytreat, purple) treatments. (F) The velocity magnitudes for stiff and soft substrates^12^. For (D) and (F), statistically significant differences are indicated (*p < 0.05, **p < 0.01, ANOVA with Benjamini-Hochberg procedure).

To validate our predictions on the impact of inhibiting oscillations on the wave speed, we next examined the changes in the wave speeds of DCs after treatment with blebbistatin, cytochalasin D, and Y27632 (which have been shown to reduce the podosome oscillations in Fig. 2D). We applied a recently developed technique, sliding time window spatiotemporal image correlation spectroscopy (twSTICS)^10^, to measure the velocity (magnitude and direction) of flowing fluorescent F-actin imaged within the cell (refer to *Models and Metho*ds). The measured velocity magnitude can be used to characterize the wave speed of podosome chemo-mechanical diffusion waves (Fig. 5D and Movie S4). By using the twSTICS to process the time series of DCs transfected with LifeAct-RFP, we quantified the wave speed for the control and cytochalasin D, blebbistatin, or Y27632 treatments on DCs (Fig. 5E and S5C-S5E). Importantly, our experiments show that the magnitudes of F-actin velocities are significantly reduced after cytochalasin D, blebbistatin, or Y27632 treatments (Fig. 5D-5E, and S5E). These observations are in agreement with our theoretical predictions (Fig. 5C). Together, these results indicate that the inhibition of podosome oscillations disrupts the chemo-mechanical diffusion waves, which also confirms the predictions of our model.

### Podosome clusters probe substrate stiffness by modulating the exchange rate between F-actin and G-actin

Next, to understand mechanosensing of podosome clusters, we investigate how podosome clusters respond to substrates with varied stiffnesses. For individual podosomes, our simulations show that higher substrate stiffness increases the steady-state protrusive force *F*_*ps*_ =*F*_*p*0_*k*_*s*_/(*k*_*s*_ + *k*_*c*_)(1 −*V*_*d*_/*V*_*pms*_), while reduces the substrate displacements *l*_1*s*_ =*F*_*ps*_/*k*_*s*_ (Fig. S6A), in line with the experimental measurements^7,12^. Microscopically, this increased protrusive force on stiffer substrates is due to the stronger and denser F-actin networks in the podosome core^3,8^. This mechano-sensitive structural difference affects the cluster dynamics because denser core actin (i.e., larger core area *A* or planar density *ρ*) on stiffer substrates requires more G-actin to assemble and releases more G-actin on disassembly (Fig. 5B). Since *μ* ∝ *Aρ*, the parameter *μ* should be proportional to substrate stiffness (*μ* ∝ *k*_*s*_), such that the effective exchange rate between F-actin and G-actin increases with substrate stiffness, i.e., *ω*_*p*_=*βμV*_*d*_/*V*_*pms*_ ∝ *k*_*s*_. Based on the analytical approximation for the propagation velocity *v*_*c*_ (Eq. 10 in *Models and Methods*), we can see that a smaller exchange rate *ω*_*p*_reduces the chemo-mechanical wave propagation speed *v*_*c*_ (Fig. 5C). Reducing the sensitivity parameter *μ* in either our continuum or discrete model leads to a lower propensity for wave-like dynamics in the podosome cluster (Fig. 5F). This is because a smaller actin exchange rate *ω*_*p*_ causes less variance (or fluctuation) in the spatial distribution of actin concentration, reducing the gradient of actin concentration and eventually decreasing the wave propagation speed *v*_*c*_ (via Eq. 10).

To validate the above predictions, we examined podosome cluster dynamics by seeding DCs on polydimethylsiloxane (PDMS) substrates with different stiffness. Using the twSTICS analysis, we found that the wave speeds are reduced on soft substrates (Fig. 5F). To further confirm that the slow wave speeds are due to small actin exchange rates rather than changes in the podosome oscillation periods, we extracted the oscillation periods of individual podosomes and found that the oscillation periods for soft and stiff substrates are indeed not significantly changed (Fig. S6B). However, the correlation coefficient decays much faster with pair distance for podosome clusters on soft substrates, indicating a smaller chemo-mechanical diffusion wavelength for soft substrates (Fig. S6C). Taken together, these experimental findings confirm our predictions that stiffer substrates can enhance the propagation of chemo-mechanical diffusion waves. Therefore, podosome clusters can serve as a mechanosensing platform by modulating the exchange rate between core F-actin and free G-actin of individual podosomes.

## Discussion and Conclusions

In this study, by developing a chemo-mechanical model and validating the predictions with experiments, we have elucidated the mechanisms of individual podosome oscillations and the wave-like dynamics of podosome clusters. Following a bottom-up approach, we first showed that two competing force generation processes of polymerization-associated protrusion at the core and myosin dynamics in the ring occur at similar rates, causing the vertical oscillations of individual podosomes. Next, our model reveals that the oscillatory growth of individual podosomes leads to release or consumption of G-actin locally, which in turn causes diffusion of G-actin and drives the wave-like coordination among the podosomes in a cluster. It is the first theoretical model, to our knowledge, that systematically illustrates how the vertical dynamics of individual podosomes are synchronized to form wave-like dynamics in podosome clusters of DCs and how the podosomes in a cluster can collectively probe mechanical cues from surroundings.

For individual podosomes, the two force-generating processes—actin polymerization for core protrusive forces and myosin dynamics for ring contractile forces—are balanced in a dynamic manner. Our model shows that the core height, ring components, and protrusion forces of podosomes all oscillate with time, in agreement with experimental results presented in this and in previous studies^7,9,10^. Our analysis reveals that the oscillatory growth of podosomes is determined by the timescale governing the core protrusion process *τ*_*p*_ and timescale for myosin turnover *τ*_*m*_. Podosomes spontaneously oscillate when these timescales are comparable, and the oscillation period can be estimated as 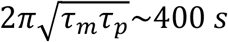. In addition to the oscillatory protrusion pattern, podosomes can exhibit monotonic growth when these timescales are not comparable, which has been validated by pharmacological treatments targeting polymerization and myosin activity. In addition, the Rho-ROCK pathway, a critical mechano-sensitive mediator closely associated with vertical oscillations of podosomes, has not been considered in previous work analyzing the dynamics of individual podosomes ^6,7,22^. Here, our model predicts that the weak mechano-sensitive feedback *Γ* due to this pathway inhibits podosome oscillations and this was validated by the experiments that inhibit this pathway (Y27632 treatment). The impacts of different pharmacological treatment on podosome growth patterns have been summarized in a phase diagram (Fig. 2C).

The oscillatory behaviors in lengths and key components, such as actin filaments and adaptors, have also been found in other protrusion types, such as invadopodia^21,23,24^ and filopodia^25,26^. Different from invadopodia and filopodia, podosomes possess a complex conical structure in which ventral actin filaments branch from the core F-actin and connect with ring adhesion. Although these complex structural components are found in individual podosomes, the dynamics of the ring components and core actin are highly synchronized and both dynamics can be mediated by myosin contractility. Therefore, the actin polymerization and myosin contractility remain the two most dominant processes governing the podosome dynamics. Furthermore, our previous model on invadopodia dynamics^21^ and other models on filopodia^25^ have also shown that their oscillatory growths are caused by interplay between actin polymerization process and myosin dynamics. Taken together, these studies suggest that the ubiquitous nature of oscillations in such nonlinear cellular protrusion systems can originate from the competing force generating processes generating compressive (actin polymerization) and tensile forces (myosin contractility) in actin networks.

Our chemo-mechanical model shows that G-actin diffusion can synchronize the vertical oscillations of individual podosomes in a cluster to form chemo-mechanical diffusion waves. Using discrete and continuum approaches, we simulate both the radial and random waves of the core F-actin observed in experiments. By evaluating the correlations between podosome dynamics in both our simulations and experiments, we found that individual podosome dynamics becomes gradually uncorrelated as their distances become larger than the characteristic wavelength, *λ*∼3 *μm*. Our model also shows that the wave propagation speed *λ*/*T* is regulated by individual podosome oscillation period, *T* and core-actin exchange rate, *ω*_*p*_. This prediction has been further validated by our experiments with pharmacological treatments to inhibit the oscillations and by varying substrate stiffness to change the core-actin exchange rate. It is also worth noting that we chose not to include the effects of dorsal filaments that connect with adjacent podosomes in our model, because the dorsal filaments mainly apply horizontal forces on the podosome core; these horizontal forces are balanced at the core, and hence, do not affect podosome dynamics.

Overall, by integrating the functions and processes of key molecular components (i.e., myosin, adaptors, G-actin, F-actin) in DCs, our model demonstrates how the interplay of these components triggers spontaneous oscillations in individual podosomes and how these spontaneous oscillations lead to wave-like patterns in clusters through diffusion. DCs can capture antigens, migrate to lymphoid tissues, and initiate the primary immune responses, a critical process in wound healing and inflammation^27,28^. Owing to the antigen-presenting and migratory capacities of DCs, DCs that present tumor antigens have become an essential target of efforts to develop therapeutic immunity against cancer^29,30^. As podosomes play a critical role in transmigration of DCs, our chemo-mechanical model can be readily adapted to understand the roles of podosome dynamics in mechanosensing and migration of DCs. Future theorical developments and computational models may build upon our chemo-mechanical model by incorporating matrix degradation and other features of the microenvironment (e.g., matrix nonlinear elasticity, matrix heterogeneity, and topographical cues), leading to a comprehensive understanding of podosome dynamics in the context of wound healing and cancer immunotherapy.

## Models and Methods

### 1. Model Formulation

The equations in the *Results* govern the spatiotemporal dynamics of a podosome cluster can be expressed as:

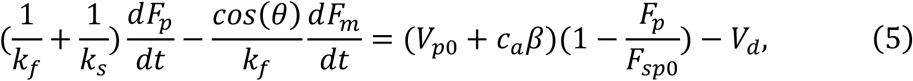

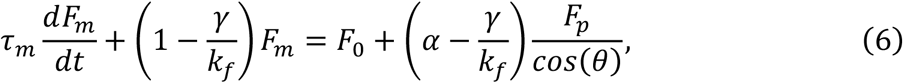

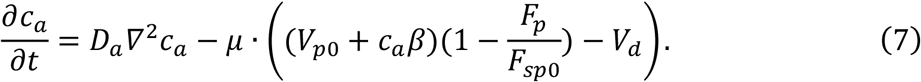

There are three variables: the protrusive force *F*_*p*_(*x,y,t*), the active myosin force *F*_*m*_(*x,y,t*) and the G-actin concentration *c*_*a*_(*x,y,t*) that describe polymerization-associated protrusion, signaling-associated myosin recruitment, and reaction-diffusion of G-actin, respectively. When the maximum polymerization speed *V*_*p*0_ + *c*_*a*_*β* is constant, Eqs.4-5 reduce to the model for individual podosome oscillations (i.e., Eqs. 1-2 without Gaussian noise). To solve Eqs. (5-7), we applied both discrete and continuum approaches using MATLAB and COMSOL packages. All the parameters used in the simulations are summarized in Table S1, and more details on the simulations can be found in SI Note 3.

### 2. Linear stability analysis

To analytically obtain the wave periods and wavelength, we performed a linear stability analysis on our chemo-mechanical model. In steady state (i.e., when the time derivatives in Eqs. 5-7 vanish), we can obtain the steady-state protrusive force *F*_*ps*_ =*F*_*sp*0_(1 −*V*_*d*_/*V*_*pms*_), myosin force 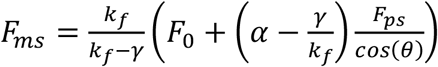 and actin concentration *c*_*as*_, where *V*_*pms* =_*V*_*p*0_ + *c*_*as*_*β* denotes the maximum polymerization speed in steady-state. Next, we applied a small perturbations, *δF*_*ps*_, *δF*_*ms*_∼ *e*^*iωt*^ to the steady-state forces, (*F*_*ps*_, *F*_*ms*_) in Eqs. 5-6, where the steady-state actin concentration *c*_*as*_ and maximum polymerization speed *V*_*pms*_ are assumed not to vary with time. Thus, we can write the eigenvalues that characterize the dynamics of the perturbed system:

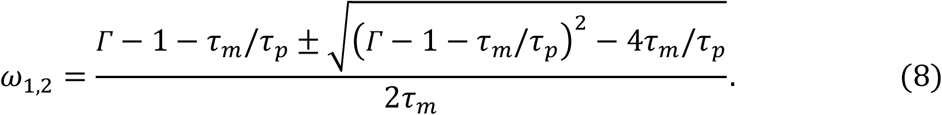

Here 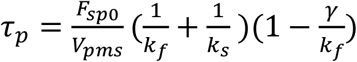 is the timescale that characterizes polymerization-associated core protrusion, *τ*_*m*_ is the timescale for myosin turnover, and 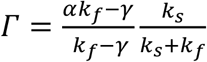 is the parameter group for signaling-associated contraction of ventral actin filaments. The eigenvalues in Eq. 8 allow us to determine the dynamics of individual podosomes: Podosomes spontaneously oscillate only when the eigenvalues’ imaginary part 𝒥_*m*_(*ω*_1,2_) ≠ 0, i.e., (*Γ* −1 −*τ*_*m*_/*τ*_*p*_)^2^< 4*τ*_*m*_/*τ*_*p*_. For oscillations with constant amplitudes, the real part *R*_*e*_(*ω*_1,2_) =0, i.e., *τ*_*m*_/*τ*_*p*_=*Γ* −1, and the oscillation periods can be estimated as 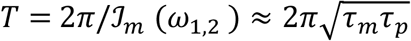 The system becomes unstable when the real part ℛ_*e*_(*ω*_1,2_) > 0, i.e., *Γ* > 1 + *τ*_*m*_/*τ*_*p*_.

Next, to obtain the wavelength for collective wave dynamics, we apply a small variation *δF*_*ps*_, *δF*_*ms*_, and *δc*_*as*_ to the reaction-diffusion Eq. 7 when it is in steady state (*F*_*ps*_, *F*_*ms*_, *c*_*as*_):

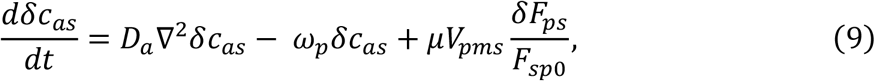

where *ω*_*p*_=*βμ*(1 −*F*_*ps*_/*F*_*sp*0_) =*βμV*_*d*_/*V*_*pms*_ is a force-mediated exchange rate between core F-actin and free G-actin, characterizing the increase of polymerization speed per nanometer decrease in core height (core F-actin disassembly releases G-actin, elevating local G-actin density and hence polymerization speed). The Eq.9 is a diffusion wave equation^31^ with the oscillatory driving force term 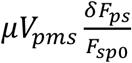 By assuming the complementary solution for Eq. 9 (i.e., setting driving force term as zero) in the form *δc*_*as*_∼ *e*^*iωt*+*iqx*^, we can get that the relation between the angular wave number *q* and angular frequency *ω*, i.e., 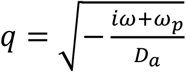 Therefore, the wavelength can be written as: 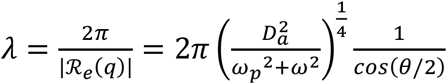 Here the phase angle is θ =π + *tan*^−1^(*ω*/*ω*_*p*_) and *ω* is assumed to be real. Furthermore, we can calculate the phase velocity *v*_*c*_ for the chemo-mechanical diffusion waves as:

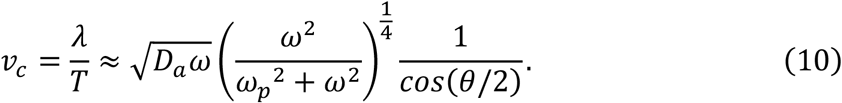

The above eigenvalues Eq.8 and the *q* −*ω* relation can be also gained by applying the small perturbation to Eqs. 5-7 simultaneously, refer to SI Note 5.

### 3. Preparation of human DCs

DCs were generated from peripheral blood mononuclear cells as described previously^32,33^. Monocytes were derived either from buffy coats or from a leukapheresis product. Plastic-adherent monocytes were cultured in RPMI 1640 medium (Life Technologies) supplemented with fetal bovine serum (FBS, Greiner Bio-one), 1 mM Ultra-glutamine (BioWhittaker), antibiotics (100 U ml−1 penicillin, 100 μg ml−1 streptomycin and 0.25 μg ml−1 amphotericin B, Gibco), IL-4 (500 U ml−1) and GM-CSF (800 U ml−1) in a humidified, 5% CO2-containing atmosphere.

### 4. DC transfection

Transient transfections were carried out with the Neon Transfection System (Life Technologies), as previously reported^10^. Briefly, DCs were washed with PBS and resuspended in 115 μl Resuspension Buffer per 0.5 × 10^6^ cells. Subsequently, cells were mixed with 6 μg DNA per 10^6^ cells per transfection and electroporated. Next, cells were quickly transferred to WillCo-dishes (WillCo Wells B.V.) with pre-warmed medium without antibiotics or serum and allowed to recover for 3 h at 37°C. For investigating the effect of substrate stiffness, cells were seeded on dishes spincoated with PDMS (Sylgard 184, Dow Chemicals; base-curing ratio 1:20 = stiff, ∼800kPa; 1:78 = soft, ∼1 kPa)^12^. Finally, the medium was replaced by a medium supplemented with 10% (v/v) FCS and antibiotics for up to 24 h. Prior to live-cell imaging, cells were washed with PBS and imaging was performed in RPMI without Phenol red. All live cell imaging was performed at 37°C.

### 5. Image analysis

All image analysis was performed using Fiji^34^ and the data was subsequently processed using Matlab (MathWorks). Prior to analysis, movies were registered, allowing translation only, using the StackReg plugin^35^ and bleach correction was performed using the “Histogram Matching” option. For amplitude ratio analysis, podosome centers were detected in the first frame of the image series with a slightly adapted ImageJ algorithm that was developed previously^36^. Briefly, before median filter processing, we first performed an unsharp mask filter (radius: 4.5 pixels, weight: 0.8) and this sequence was repeated twice after which a background subtraction was performed. Podosome centers were subsequently determined using the ImageJ maxima finder (prominence: 1000). Fluorescence intensity was measured in a circular ROI with a radius (∼0.5 μm) over the course of the movie.

### 6. Extracting oscillation amplitudes, periods, and correlation coefficient

To extract the oscillation amplitude and frequency, the measured fluorescence intensity dynamics were converted from the time domain to frequency domain using Fast Fourier transformation (FFT). The largest four peaks in the spectrum were chosen to calculate the amplitudes and frequencies of the podosome dynamics (Fig. S4). The amplitude ratio was calculated as the ratio of the averaged amplitudes and the mean intensity. Next, to estimate the spatial correlation of two podosomes in a cluster, Pearson correlation coefficients were calculated for the time series of fluorescently tagged LifeAct over a 25 min time window. By denoting the LifeAct intensity dynamics of two podosomes as vectors *A* and *B* of length *N*, the correlation coefficient is written as:

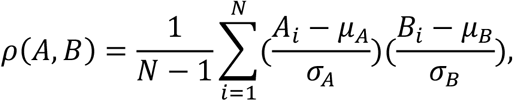

where *μ*_*A*_ and *σ*_*A*_ are the mean and standard deviation of *A*, respectively, and *μ*_*B*_ and *σ*_*B*_ are the mean and standard deviation of *B*, respectively. The correlation has a value between −1 and 1, which correspond to in phase and out-of-phase correlation, respectively. The *corrcoef* function in Matlab (MathWorks) is used to calculate the correlation coefficient.

### 7. Time window STICS analysis

We performed Spatio-temporal Image Correlation Spectroscopy (STICS)^37^ with a short time window iterated in single frame shifts on a CLSM and Airyscan time series of fluorescently tagged Lifeact, acquired with a 15 s time lag between frames. First, we applied a Fourier immobile filter in time to each pixel stack in the entire image series to remove the lowest frequency (e.g. static) components^37^. Next, we divided each image into 16×16 pixels ROIs (2.24 μm x 2.24 μm for the CLSM and 0.64 μm x 0.64 μm for the Airyscan) and shifted adjacent ROIs four pixels in the horizontal and vertical directions to map the entire field of view with oversampling in space. We then divided the time series into overlapping 10 frames sized TOIs (2.5 min) and shifted adjacent TOIs one frame for each STICS analysis to cover the entire image series with oversampling in time. We then calculated space-time correlation functions for each ROI/TOI and fit for time lags up to τ = 8 to measure vectors (magnitude and direction) of the flow from the translation of the correlation peak. Due to the spatial and temporal oversampling (75% common overlap in space between adjacent ROIs and 90% common overlap in time between sequential TOIs) we expect neighboring vectors to correlate in magnitude and direction for real flows. Further details about the detection and elimination of noise vectors caused by random fits to spurious background peaks and the twSTICS method in general have been previously described^10^. The movies were collected from at least 2 experiments for each condition.

### 8. Statistical analysis

Statistical analyses were performed using Matlab (MathWorks), and data were compared using the one-way ANOVA with Benjamini-Hochberg Procedure or Student test as indicated in figure legends. The error bars in Fig. 4G, S4D, and S6 represent standard deviation (SD), and the sample sizes and p values are given in figure legends.

## 9. Data and code availability

Experimental datasets and codes are available upon request from the corresponding author.

## Acknowledgements

This work was supported by National Cancer Institute awards R01CA232256 and U54CA261694; National Institute of Biomedical Imaging and Bioengineering awards R01EB017753 and R01EB030876; NSF Center for Engineering Mechanobiology Grant CMMI-154857; NSF Grants MRSEC/DMR-1720530 and DMS-1953572. Experimental work was supported by Intramural funding from Radboud University Medical Center to AC.

## Author Contributions

Z.G. and V.B.S. conceived the chemo-mechanical model and carried out the theoretical and numerical analyses. A.C. and K.D. conceived the experiments. K.D. performed the experiments. Z. G and K.D. analyzed the experimental data. Z.G., K.D., A.C., and V.B.S. wrote the paper.

## Declaration of Interests

The authors declare no conflict of interest.

## Supplementary Information

Document S1: Figures S1-S9, Table S1, and Notes S1-S4

Movie S1: Podosome cluster in a representative DC with Lifeact-RFP and vinculin-GFP transfected showing oscillations of individual podosomes and random waves in podosome cluster.

Movie S2: Simulated podosome cluster with the discrete approach showing the propagation of radial and random waves

Movie S3: Simulated podosome cluster with the continuum approach showing the propagation of radial and random waves

Movie S4: Representative same DC before and after adding cytochalasin D. DCs are transfected with LifeAct-RFP using confocal microscopy with 15 s frame intervals.

## Notes

### Competing Interest Statement

The authors have declared no competing interest.

