## Supplementary figures, table and texts for "Chemo-mechanical Diffusion Waves Orchestrate Collective Dynamics of Immune Cell Podosomes"

**for**

**Supplementary Figures and Tables**

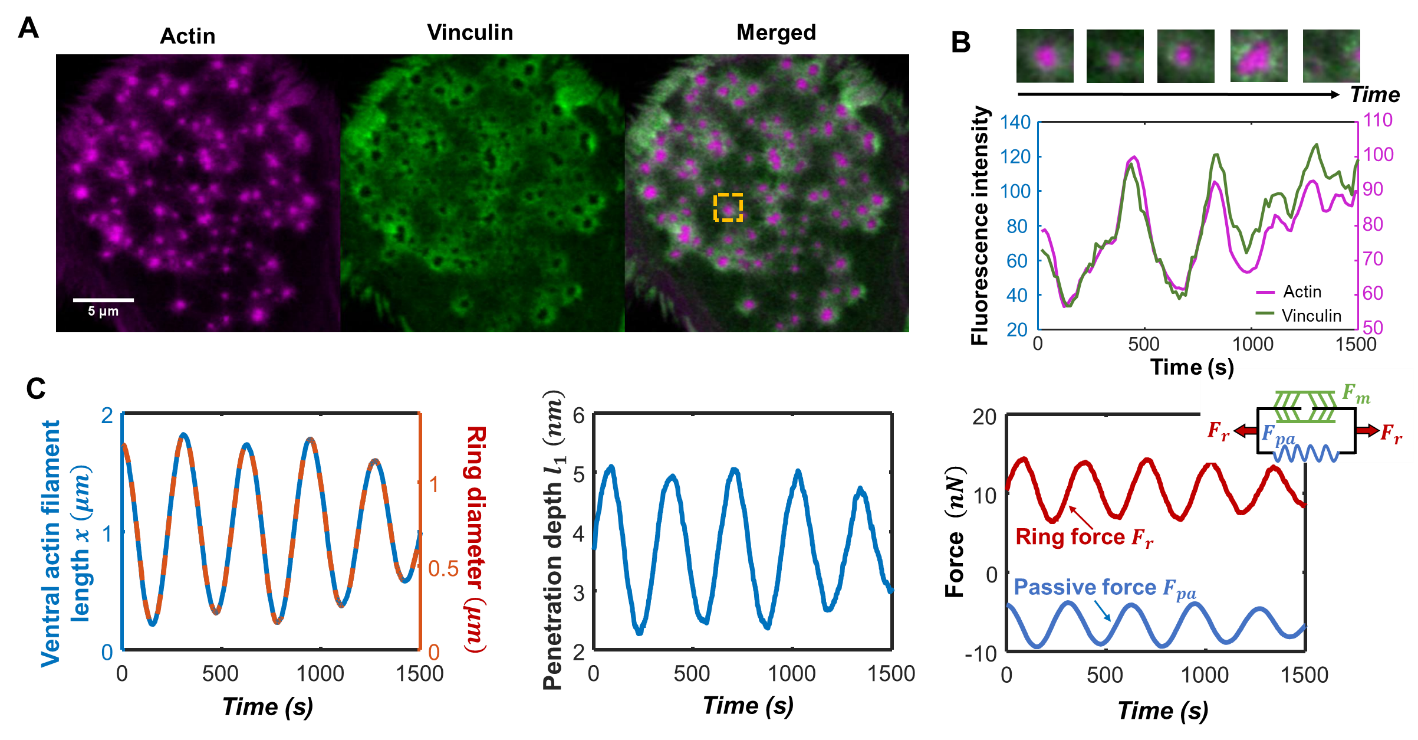

**Fig S1.** **Oscillatory behaviors in podosome core and ring.** (A) Images of a DC stained for actin (magenta) and vinculin (green). (B) The experimentally measured fluorescence intensity of actin (magenta) and vinculin (green) for a representative podosome plotted versus time. The insets shown the time series for the representative podosome. (C) The simulated (left panel) ventral actin filament length (blue line), ring diameter (red line), (middle panel) substrate displacement, and (right panel) forces plotted with time.

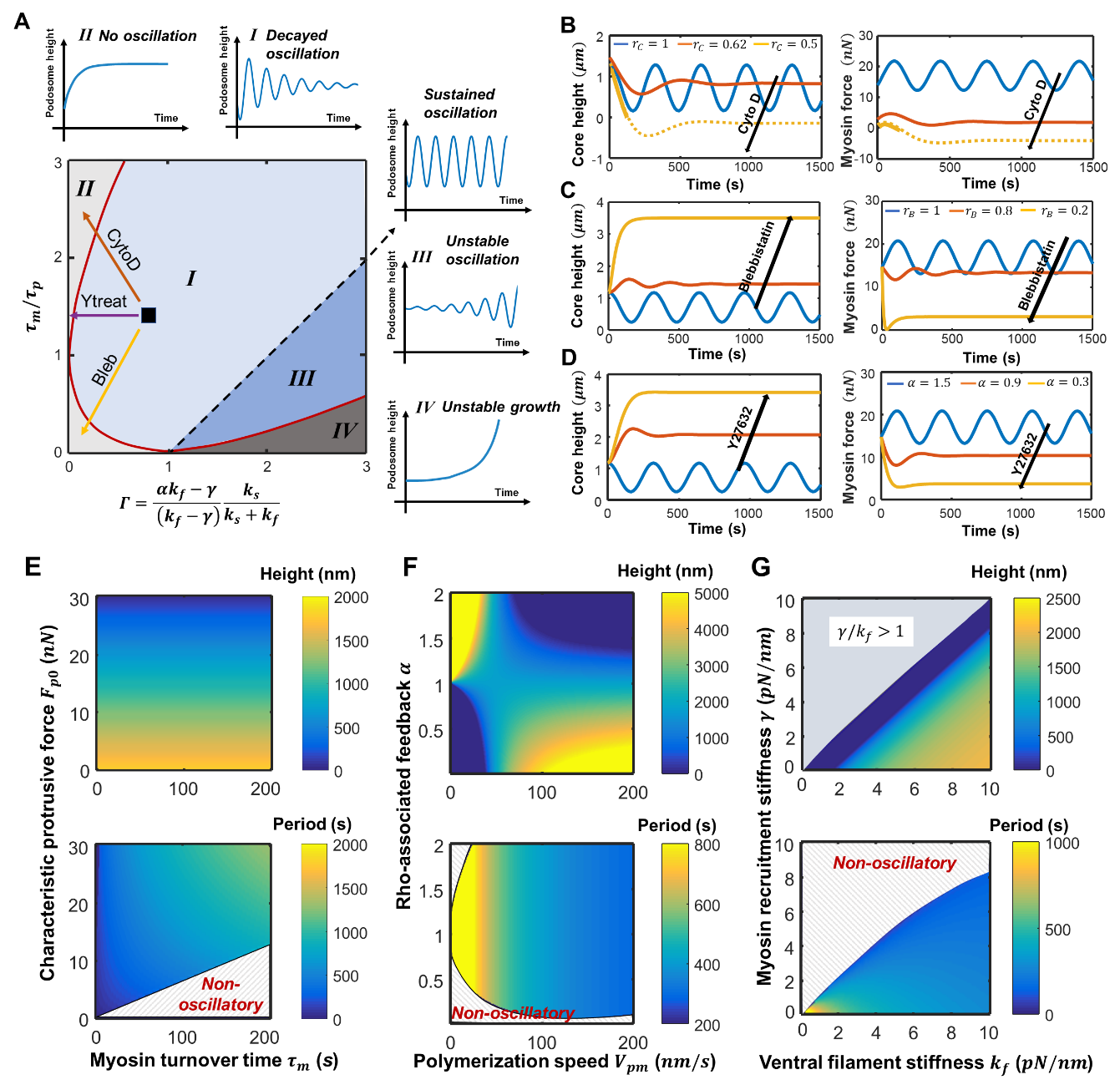

**Fig S2. Phase diagram and parameter sensitivity analysis.** (A) Phase diagram showing the different types of protrusion patterns based on timescale ratio $\tau_{m}/\tau_{p}$ and the feedback parameter $\Gamma$. The arrows indicate the influence of cytochalasin D (CytoD, red), blebbistatin (Bleb, yellow), and Y27632 (Ytreat, purple) treatments on the dynamics. (B-D) The simulated (left panels) core height and (right panels) myosin force plotted with time for (B) cytochalasin D, (C) blebbistatin, and (D) Y27632 treatments. (E-G) The simulated (Top panels) core height and (Bottom panels) oscillation period plotted for (E) myosin turnover timescale and maximum protrusion force, (F) polymerization speed and Rho-associated feedback, and (G) ventral filament stiffness and effective stiffness for myosin contraction. The non-oscillatory regimes are marked with grey strips.

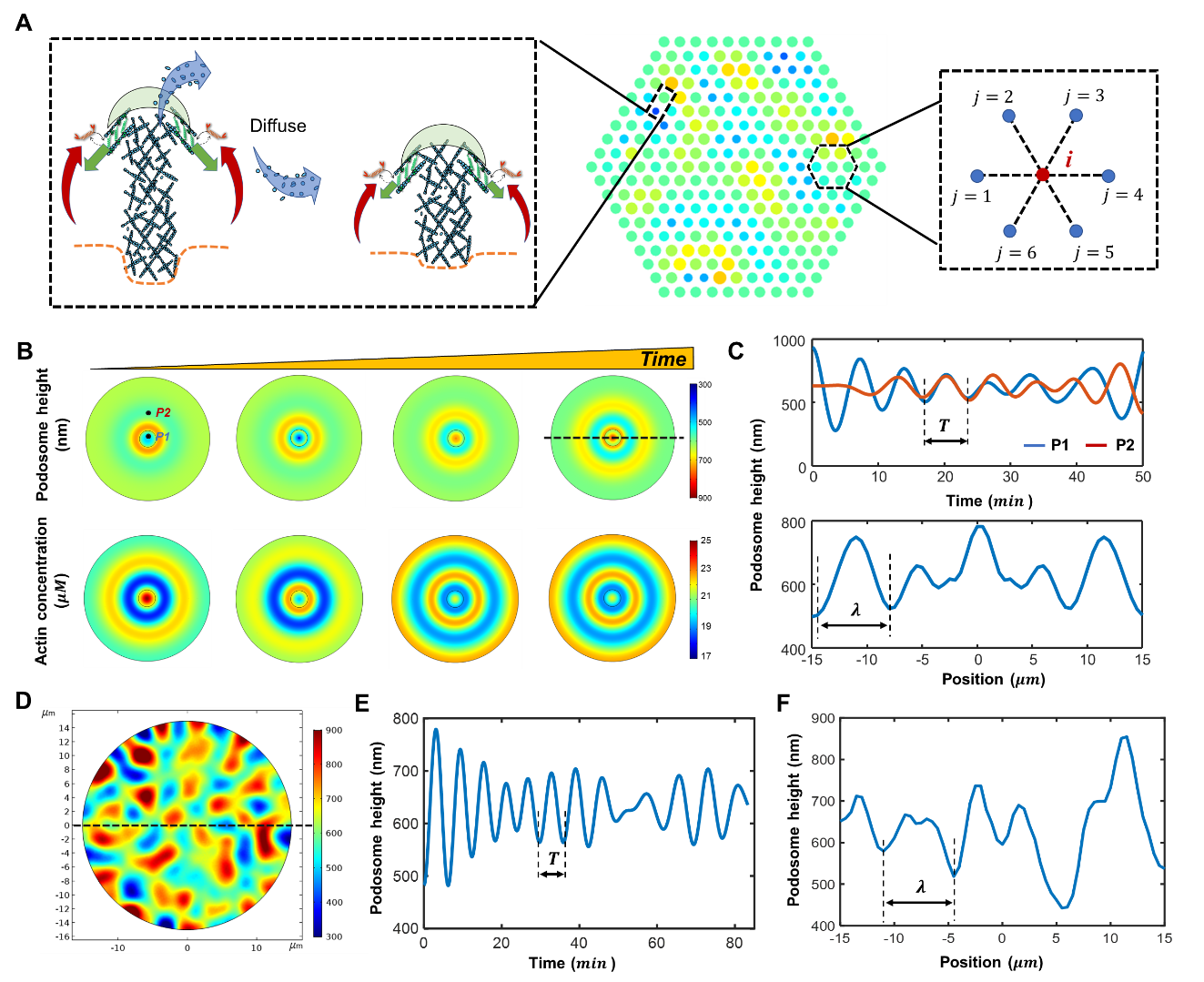

**Fig S3.** **The simulated spatiotemporal dynamics for radial and random waves.** (A) The discrete model showing the random wave patterns of podosome height. Individual podosomes are at a fixed separation $d_{0}=1.5 \mu m$, and the dynamics of different podosomes are correlated through actin diffusion. (Left inset) Schematic for two neighboring podosomes in the discrete model. (Right inset) The podosome $i$ and its neighboring podosomes *j*=1, 2, …,6. (B) (Top panels) The simulated podosome core heights and (Bottom panels) actin concentration in the cluster plotted for different time. (C) (Top panel) The podosome height plotted with time for two representative points marked in panel (B). (Bottom panel) The podosome height plotted for different positions in the dashed cutting line in (B). (D) Simulated heights of a podosome cluster showing random waves. (E) The podosome height plotted with time for a representative point. (F) The podosome height plotted for different positions in the dashed cut line in (D). The wave periods $T$ and wavelengths $\lambda$ are labeled in (C), (E), and (F).

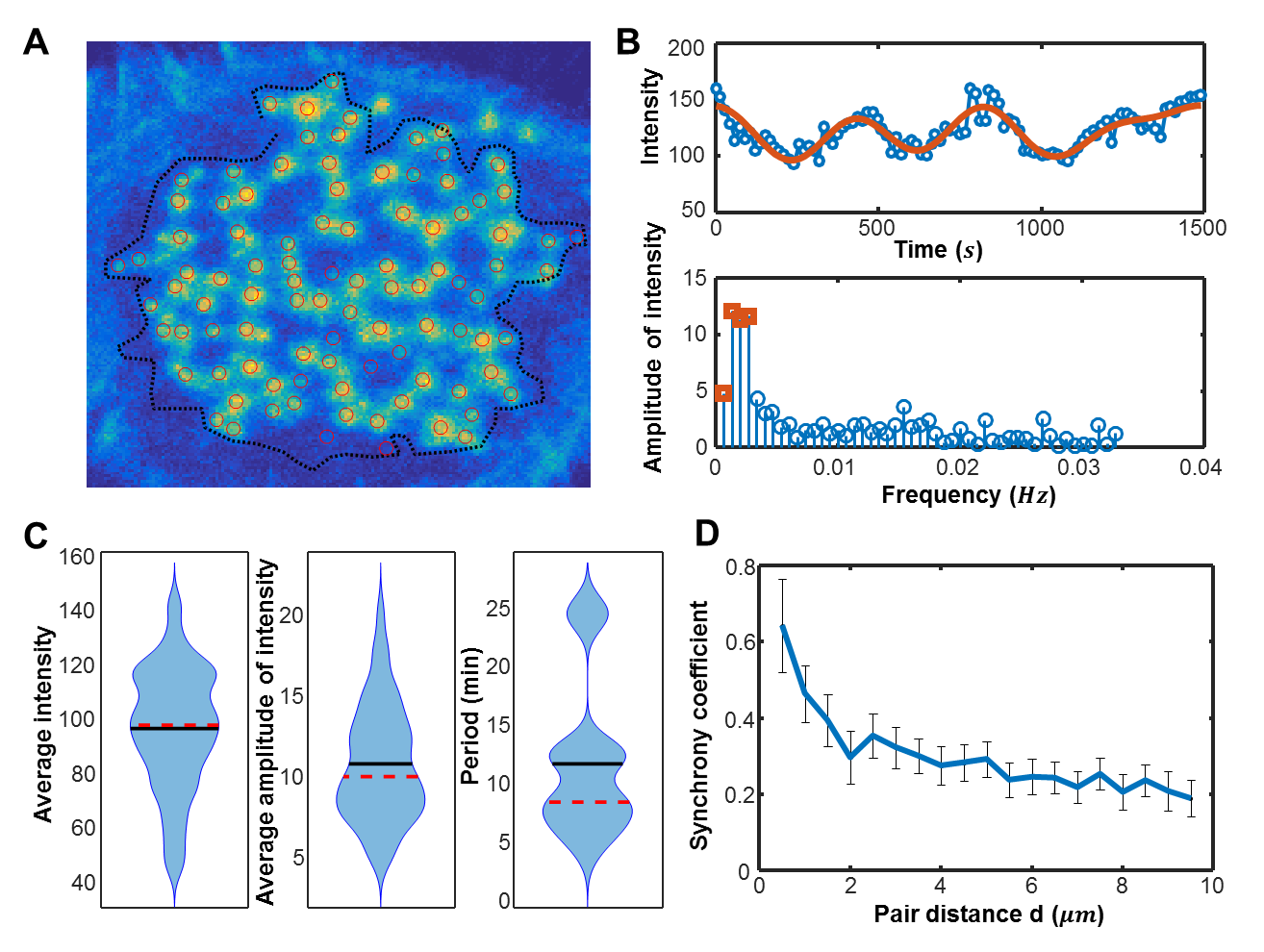

**Fig S4.** **Extraction of podosome oscillation periods and the synchrony coefficient in a podosome cluster.** (A) A representative LifeAct-RFP transfected DC. The red circles locate the podosome positions extracted using ImageJ, and the dashed line indicate the region of interest. (B) (Top panel) The fluorescence intensity of a representative podosome in (A) plotted with time. (Bottom panel) The frequency spectrum showing the amplitudes of intensity extracted by Fast Fourier Transformation (FFT) as a function of frequency. Blue dots and line represent the experimentally measured dynamics, and red line in the top panel indicate the time-domain dynamics transformed from the largest four amplitude peaks in the spectrum (i.e., red markers in bottom panel) using FFT. (C) The violin plots for (left panel) average fluorescence intensity, (middle panel) averaged amplitude, and (right panel) period for the representative DC shown in (A). Podosome number n=89. (D) The synchrony coefficient plotted as a function of pair distance for the representative DC shown in (A). Podosome pair number n=3916.

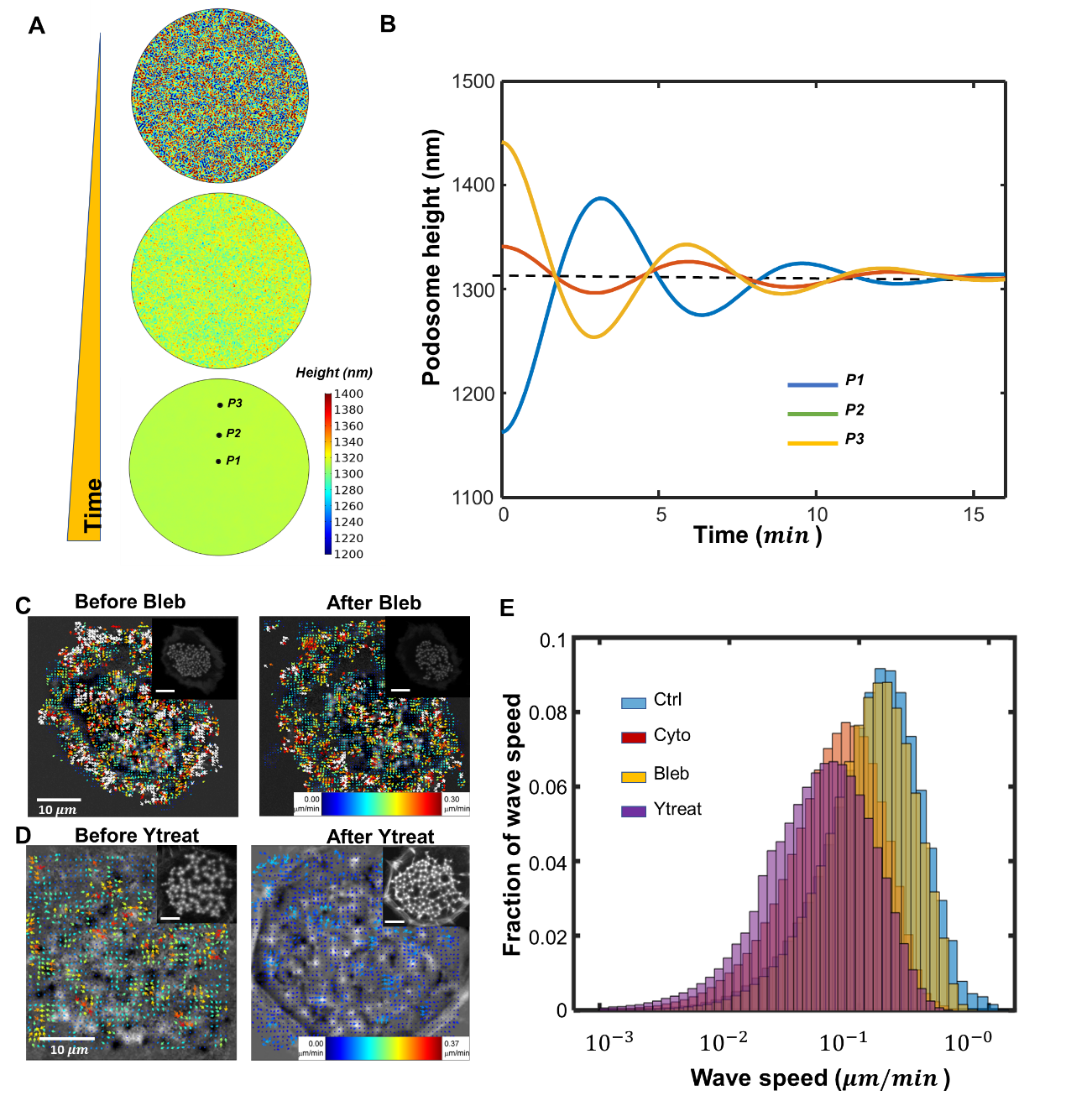

**Fig S5.** **Disrupted diffusion waves after the pharmacological treaments.** (A) The simulated podosome core heights in the cluster plotted for different instances of time. (B) The ventral actin filament length plotted versus time for three representative points marked in (A). (C-D) Representative DC (left panel) before and (right panel) after adding (C) Blebbistain and (D) Y27632. DCs are transfected with LifeAct-GFP using confocal microscopy at 15 s intervals. Time series subject to twSTICS analysis were plotted as vector maps; the insets show the corresponding DCs without twSTICS analysis. The arrows indicate flow directions, and both the size and color denote the ﬂow magnitude. (E) Fractions of wave speeds for control (Ctrl, blue), cytochalasin D (Cyto, red), Blebbistatin (Bleb, yellow), and Y27632 (Ytreat, purple) treatments.

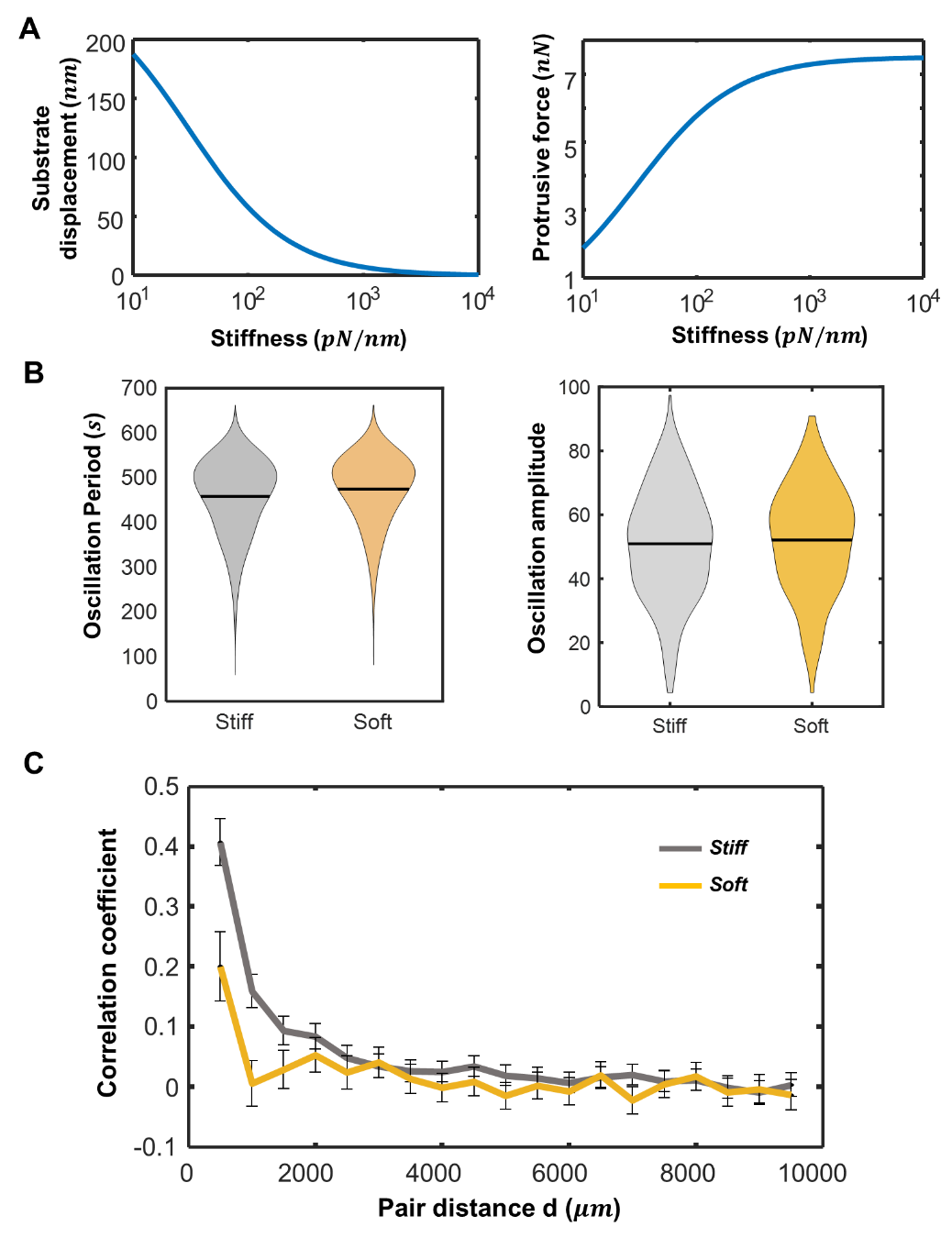

**Fig S6.** **Mechanosensing in podosome clusters.** (A) The simulated (left panel) substrate displacement and (right panel) protrusive force plotted for different levels of stiffness. (B) The extracted (left panel) oscillation periods and (right panel) amplitudes plotted for stiff and soft substrates. No significant difference observed between the two plots. (C) The correlation coefficient plotted as a function of pair distance for stiff and soft substrates.

**Table S1. General model parameters.**

| **Parameters** | **Meaning** | **Value** | **Reference** |
| --- | --- | --- | --- |
| $F_{p0}$ | Characteristic protrusive force | 20 nN | ^1,2^ |
| $V_{p0}$ | Initial polymerization speed | 80 nm/s | ^3,4^ |
| $V_{d}$ | Depolymerization speed | 50 nm/s | Adjust based on ^5,6^ |
| $\beta$ | Sensitivity for polymerization speed for a $\mu M$ change in G-actin concentration | 0.5 $nm/s/\mu M$ | Adjusted based on ^7^ |
| $k_{s}$ | Substrate stiffness | 10-2000 $kPa$ | ^3,8^ |
| $D_{a}$ | Diffusion coefficient | 0.03 $\mu m^{2}/s$ | Fitting parameter, refer to SI Note 3. |
| $\alpha$ | Rho-ROCK feedback parameter | 1.5 | Adjusted based on ^9^ and linear stability analysis, refer to SI Note 3. |
| $\gamma$ | Effective stiffness for myosin contraction | 0.2 $pN/nm$ |  |
| $k_{c}, k_{f}$ | Stiffness for core F-actins and ventral F-actins | 10 $pN/nm$  3.5 $pN/nm$ |  |
| $c_{as}$ | Steady state G-actin concentration | 20 $\mu M$ | ^7,10^ |
| $\chi_{p}, \chi_{m}$ | Gaussian noise for actin polymerization and myosin recruitment | $\sigma\left( \chi_{p} \right)=0.1V_{p0}$  $\sigma\left( \chi_{m} \right)=0.1F_{p0}$ | Adjusted base on ^4^, refer to SI Note 3. |
| $\theta$ | Angle between the ring ventral F-actins and the core | $\pi/4$ | Estimated in this work |
| $\tau_{m}$ | Characteristic time for myosin turnover | 35 s | Adjusted based on ^11,12^ |
| $\mu$ | Local actin concentration change for the growth of podosome height by a micron | 0.06 $\mu M/\mu m$ | Fitting parameter, refer to SI Note 3. |
| $F_{M0}$ | Initial myosin force | 1000 pN |  |
| $x_{0}$ | Initial length of ventral actin filaments | 2.8 $\mu m$ |  |
| $d_{0}$ | Neighbouring podosome distance in the discrete model | 1.5 $\mu m$ | ^13,14^ |
| $R_{0}$ | Podosome cluster radius | $15 \mu m$ | Measured in experiments |

**SI Note S1. Molecular basis for mechano-sensitive recruitment of myosin**

The tensile forces sustained by the ventral actin filaments (VAFs) consist of the active myosin contractility and the passive elastic force ^9^. The dynamics of active myosin contractility is determined by the processes of myosin recruitment (binding to VAFs) and turnover (unbinding from VAFs). Assuming that there is a large cytoplasmic pool of myosin, of which the total myosin number is much larger than the bound myosin number. (i.e.*,* $m_{0}\gg m_{b}$), the dynamics by bound myosin is:

$$\frac{dm_{b}}{dt}\approx k_{on}^{*}m_{0}-k_{off}m_{b}+\xi_{m}(t). (S1)$$

Here $k_{on}^{*}$ and $k_{off}$ are the rates for myosin recruitment and turnover, respectively, and the $\xi_{m}(t)$ is a Gaussian noise term to represent the fluctuations in myosin number during myosin dynamics. As discussed in the *Results* section, the mechano-sensitive biochemical pathways (Rho-ROCK signaling) increases the myosin recruitment, while a large VAF displacement reduces the myosin recruitment. Therefore, we can write the effective binding rate of myosin as:

$$k_{on}^{*}=k_{on}^{0}+\frac{\alpha_{0}k_{on}^{0}}{f_{b}}F_{r}-\gamma_{0}k_{on}^{0}\frac{\left( x-x_{0} \right)}{x_{0}}. (S2)$$

Here $k_{on}^{0}$ is the initial myosin binding rate (i.e., without signaling), while $\alpha_{0}$ and $\gamma_{0}$ are non-dimensional parameters characterizing the impact on myosin recruitment from Rho-ROCK pathway and VAF displacement, respectively. The active myosin force can be further calculated as $F_{m}=m_{b}f_{b}$, where $f_{b}$ is the characteristic force per myosin. By combining all the above equations, we obtain:

$$\tau_{m}\frac{dF_{m}}{dt}+F_{m}=F_{m0}-\gamma\left( x-x_{0} \right)+\alpha F_{r}+\chi_{m}\left( t \right). (S3)$$

Here $\tau_{m}=1/k_{off}$ is the characteristic time for myosin turnover, and $\alpha=\alpha_{0}k_{on}^{0}m_{0}/k_{off}$ and $\gamma=\gamma_{0}k_{on}^{0}f_{b}m_{0}/x_{0}/k_{off}$ are two coefficients characterizing the effects of Rho-ROCK and VAF displacement on myosin recruitment, respectively. $\chi_{m}\left( t \right)=f_{b}\xi_{m}(t)/k_{off}$ represents the force fluctuation in myosin dynamics, and $F_{m0}=m_{0}f_{b}k_{on}^{0}/k_{off}$ is the myosin force equilibrium constant when signaling feedback is absent.

**SI Note S2. Model predicts individual podosome dynamics after different pharmacological treatments**

First, we studied how the inhibition of myosin contractility using Y27632 and blebbistatin (Bleb) can affect podosome dynamics. To study the influence of Rho-ROCK pathway on the podosome, we can simply reduce the signaling-associated parameter $\alpha$. In the phase diagram (purple arrow in Fig.2C and S2A), the reduced feedback parameter $\Gamma=\frac{\alpha k_{f}-\gamma}{\left( k_{f}-\gamma\right)}\frac{k_{s}}{k_{s}+k_{f}}$ leads to the non-oscillatory behaviors. By reducing the parameter $\alpha$, our simulations indeed show that the oscillatory growth of podosomes are inhibited (Fig. S2D), in agreement with our experiments (Fig. 2D). Next, to study the influence of Bleb on podosome dynamics, we consider the increase of all the parameters that are associated with myosin unbinding (with binding rate $k_{off}$ in Eq. S2). Hence, Bleb treatment decreases the myosin turnover time $\tau_{m}=1/k_{off}$ and the signaling-associated parameter $\Gamma$, corresponding to the yellow arrow in Fig. 2C and S2A. Here we defined parameter ratio $r_{m}$ ($r_{m}<1$) that is multiplied to myosin-associated parameters, i.e., $\tau_{m}$, $F_{m0}$, $\alpha$, and $\gamma$ in our model. Similar to Y27632 treatments, we found that the oscillations in the core and ring are inhibited after treatments (i.e., reduced $r_{m}$ in Fig. S2C), in agreement with our experiments (Fig. 2D). In addition to the changes in oscillatory patterns, our simulations show that the time-averaged core heights increase after either the Y27632 or Bleb treatment (Fig.S2), as the VAF and myosin apply less contractility to restrict the core growth. Interestingly, our previous work has also shown the increased fluorescence intensity of F-actins after Bleb treatments, which again validates our model predictions.

We next assessed how the inhibition of actin polymerization using cytochalasin D (Cyto D) affects podosome dynamics. The disruption of F-actins not only influences the actin networks assembly in the core but also affects the myosin contractility at the ring, since less myosin motors are recruited to the VAFs after F-actin disruption. In our model, both the polymerization speed $V_{pm}$, VAFs’ passive stiffness $k_{f}$, the signaling-associated parameters $\alpha$ are decreased when the F-actin are disrupted by Cyto D. Hence, the core protrusion timescale $\tau_{p}=\frac{F_{sp0}}{V_{pms}}(\frac{1}{k_{f}}+\frac{1}{k_{s}})(1-\frac{\gamma}{k_{f}})$increases and signaling feedback groups $\Gamma$ decreases, and podosomes undergo non-oscillatory growth based on our phase diagram (orange arrow in Fig. 2C). To further quantify the inhibition effects of Cyto D, we defined a parameter ratio $r_{c}$ ($r_{c}<1$) that is multiplied to the polymerization speed $V_{p}$, VAFs’ passive stiffness $k_{f}$, the signaling-associated parameters $\alpha$ in our model (Fig.S2). We found that impaired F-actin inhibits the oscillatory growth in the core height (Fig. S2B). In addition to the changes in oscillatory patterns, our simulations show that the Cyto D treatment significantly reduces the core heights and ring forces, which subsequently inhibits force-mediated adaptors, i.e., vinculin and paxillin. Interestingly, our previous work has also shown that fluorescence intensity of F-actin, vinculin, and paxillin are significantly reduced after Cyto D treatments, in agreement with our simulations.

**SI Note S3. Parameter justification for the model**

**Timescales in podosome clusters:** The timescale $\tau_{m}$ for myosin turnover is chosen as $40 s$, in agreement with previous work ^15,16^. Note that the timescale $\tau_{m}$ for myosin turnover here does not necessarily describe the single myosin turnover dynamics (which happens much faster at ~1 s ), as we do not distinguish whether changes of the myosin kinetics are results of alterations in myosin turnover, changes of available cortical binding sites, or intracellular flow of myosin. The protrusion timescale for podosome core is calculated as $\tau_{p}=\frac{F_{sp0}}{V_{pms}}(\frac{1}{k_{f}}+\frac{1}{k_{s}})(1-\frac{\gamma}{k_{f}}) \sim60 s$ based on parameter values in Table S1, and both larger substrate stiffness $k_{s}$ and larger polymerization speed $V_{pms}$ reduce the protrusion timescale. When the protrusion timescale and myosin turnover timescale is comparable ($\tau_{m}/\tau_{p} \sim1$), podosomes begin to oscillate and the estimated oscillation period $2\pi\sqrt{\tau_{m}\tau_{p}}$ is around 400 s, in line with our experimental measurement (Fig. 4). Meanwhile, we can also estimate the diffusion time between neighboring podosomes in our discrete model, which is around ${d_{0}^{2}}/{6D_{a}}\sim10 s$; it is much smaller than the podosome oscillation period.

**Forces and stiffnesses in the podosome system:** In our podosome growth model, the protrusive force $F_{p}$ generated by the core actin at steady state is mainly determined by the ratio of depolymerization and polymerization speed $V_{d}/V_{pms}$ and the ratio between substrate stiffness and core stiffness $k_{c}/k_{s}$, i.e., $F_{ps}=F_{p0}k_{s}/(k_{c}+k_{s} )\left( 1-V_{d}/V_{pms} \right)$, and its magnitude is around 10 $nN$. This force magnitude also agrees with the previous measurement using the protrusive force microscopy as well as previous theoretical modelling ^1,17^. Meanwhile, the protrusive force is balanced by the forces sustained by the ring and VAFs, which consists of the contractile force $F_{m}\approx18 nN$ produced by bound myosin and the passive compressive force of VAFs $F_{pa}\approx-8 nN$ (Fig. S1). Note that, for simplicity, we assume the linear elastic materials for the substrate, as the deformations are relatively small. Although nonlinear elasticity can be easily adapted into our podosome growth model, the oscillatory patterns will not be affected by the different constitutive laws for substrates.

**Mechano-sensitive signaling feedback:** The feedback parameter group $\Gamma=\frac{\alpha k_{f}-\gamma}{\left( k_{f}-\gamma\right)}\frac{k_{s}}{k_{s}+k_{f}}$ contains mainly three parameters: $\alpha$, $k_{f}$, and $\gamma$. The parameter $\alpha$ and $\gamma$ characterize the effects of Rho-ROCK and VAF displacement on active forces (myosin contractility), while $k_{f}$ is the stiffness that characterize the VAF displacement on passive force. As the myosin contractility at steady state $F_{ms}=\frac{k_{f}}{k_{f}-\gamma}(F_{0}+\left( \alpha-\frac{\gamma}{k_{f}} \right)\frac{F_{ps}}{\cos\left( \theta\right)})$ is always positive, we have the condition $\gamma/k_{f} <1$. Besides, as the feedback groups $\Gamma=\frac{\alpha k_{f}-\gamma}{\left( k_{f}-\gamma\right)}\frac{k_{s}}{k_{s}+k_{f}}$ is positive, we have the condition $\alpha>\gamma/k_{f}$. Here, in our model, we have $\gamma/k_{f} \approx0.06$, indicating that the influence of ventral actin filament length change on the active force is negligible compared to its influence on the passive force. The values of these three parameters ($\alpha$, $k_{f}$, and $\gamma$) are adjusted based on our previous publications ^4,9^ to fit for the experimental data.

**Gaussian noise:** Two Gaussian noise terms $\chi_{p}\left( t \right)$ and $\chi_{m}\left( t \right)$ were applied to account for the random fluctuation in actin polymerization and the myosin recruitment dynamics. The Gaussian noises are generated by MATLAB *randn* function $\chi_{m}\left( t \right), \chi_{p}\left( t \right) \sim N(0,\sigma)$, where their standard deviation are estimated around 1/100 of the characteristic protrusive force ($\sigma\left( \chi_{m} \right)=0.01F_{p0}\sim1000 pN$) and initial polymerization speed ($\sigma\left( \chi_{p} \right)=0.01V_{p0}\sim1 nm/s$), respectively. In the oscillation regime I (Fig. 2C and S2A), the podosome protrusion system works as a stochastic amplifier, amplifying the small noise signal into a large oscillation in protrusion length. The random fluctuation applied here sustains the oscillation for podosomes in regime I. Note that the noise added to the system does not change the steady-state lengths significantly. In addition, we want to point out that these environmental noises can lead to time-dependent variations in length even for monotonically growing protrusions, making it difficult to differentiate the oscillatory and non-oscillatory behaviours in experiments. Therefore, here we used the amplitude ratio, defined as $r_{a}=A_{osc}/\bar{R}$, to quantify the intensity of oscillation in podosome dynamics.

**Sensitivity analysis:** Heat maps for the podosome core height at steady state and oscillation periods (non-oscillatory regimes are marked with grey strips) were plotted as a function of different key parameters (Fig. S2E-G). In our simulations, a larger characteristic protrusive force $F_{p0}$ reduces the podosome steady-state height while increases the oscillation periods, and myosin turnover timescale $\tau_{m}$ does not affect the podosome heights at steady state but increases the podosome oscillation periods (Fig. S2E). A smaller Rho-associated feedback parameter $\alpha$ reduces the myosin contractility at the VAFs, leading to larger podosome heights. Meanwhile, a larger polymerization speed $V_{pm}$ can increase the podosome core heights when Rho-associated feedbacks is small ($\alpha$<1). However, for a larger Rho-associated feedback parameter ($\alpha$>1), larger polymerization can also reduce the podosome heights (Fig. S2F). This negative regulation effect is due to that the large protrusive forces generated by large polymerization increases the ring forces, triggering more myosin recruitment through Rho-ROCK pathway, which eventually constrain the podosome growth more. Furthermore, a larger VAF stiffness increases the podosome heights, as it becomes more difficult to get compressed for VAFs and constrain the height growth, while a larger myosin recruitment stiffness reduces the heights as more myosin will be recruited to exert contraction (Fig. S2G). The regulation effects of key components are further examined using different inhibitor treatments (Cytochalasin D, Y27632, and Blebbistatin), which can also be found in the heat maps. For Bleb or Y27632 treatments, the decrease of parameter, including $\alpha$ and $\gamma$, increases the podosome height and cause podosomes to undergo non-oscillatory dynamics (Fig. S2F). For Cyto D treatment, it disrupts the polymerization in both the core actin and ring VAFs (i.e., reduce $V_{pm}$, $\alpha$, and $k_{f}$), podosome heights would reduce after the treatments and the oscillations are inhibited (Fig. S2F-G).

**SI Note S4. Discrete and continuum approaches for solving the equations**

To solve the ordinary differential Eqs. 4-5 and one diffusion Eq. 6, we applied the both discrete and continuum approaches. In our discrete model, we first generate a triangular lattice for podosome cluster, such that each node represents an individual podosome. Each podosome in the cluster is governed by the polymerization-associated protrusion (Eq. 4) and signaling-associated myosin recruitment (Eq. 5). The maximum polymerization speed in Eq.6 is regulated by the actin concentration, i.e., $V_{p0}+c_{a}\beta$, which can freely diffuse in the podosome cluster. As the distance between neighboring podosomes is the same (distance $d_{0}$), we can calculate the divergence of G-actin concentration gradient at the $i$ th podosome as:

$$\nabla^{2}c_{a}^{i,0}=\sum_{j=1}^{3} \frac{c_{a}^{i,j}+c_{a}^{i,j+3}-2c_{a}^{i,0}}{d_{0}^{2}}, j=1,2,3. (S4)$$

Here the superscript $c_{a}^{i,j}$ indicates the $j$th neighboring podosomes for the podosome $i$ (Fig. S3A), and we use $c_{a}^{i,0}$ to denote the actin concentration at the $i$ th podosome. For a podosome that is not in boundary, it has six neighboring podosomes ($j=1, 2, 3,..,6$). For those podosomes that are in the boundary, we set the gradient $\nabla c_{a}^{i}=0$ for the assumption of no boundary flow. Furthermore, we can generalize the discrete model with a continuum approach, where we treated the podosome height in the cluster as a continuous variable to indicate its spatial average. For this continuum approach, we can solve the differential Eqs. 5-7 using the Mathematics Module in COMSOL.

**SI Note S5. Analytical approximation for diffusion waves**

We can further interpret the model from the analytical approximation using the linear stability analysis. In steady state, we obtain the steady-state protrusive force $F_{ps}=F_{sp0}\left( 1-V_{d}/V_{pms} \right)$, steady-state myosin force $F_{ms}=\frac{k_{f}}{k_{f}-\gamma}(F_{0}+\left( \alpha-\frac{\gamma}{k_{f}} \right)\frac{F_{ps}}{\cos\left( \theta\right)})$ and steady-state actin concentration $c_{as}$. Note here $V_{pms}=V_{p0}+c_{as}\beta$ denotes the maximum polymerization speed in steady-state. Next, we applied a small perturbation of $\delta F_{ps}$, $\delta F_{ms}$, $\delta c_{as}$ ~ $e^{i\omega t+iqx}$ to the steady state ($F_{ps}$, $F_{ms}$, $c_{as}$) of Eqs. 5-7 simultaneously and obtained the relation between angular wavenumber $q$ and angular frequency $\omega$:

$$-\omega_{p}\left( 1+i\omega\tau_{m} \right)+(D_{a}q^{2}+i\omega+\omega_{p})(\left( 1+i\omega\tau_{m} \right)\left( 1+i\omega\tau_{p} \right)-i\omega\tau_{p}\Gamma)=0, \left( S5 \right)$$

Here $\tau_{p}=\frac{F_{sp0}}{V_{pms}k_{s}}(\frac{k_{s}}{k_{f}}+1)(1-\frac{\gamma}{k_{f}})$ is the timescale that characterizes polymerization-associated core protrusion, $\tau_{m}$ is the timescale for myosin turnover, and $\Gamma=\frac{\alpha k_{f}-\gamma}{\left( k_{f}-\gamma\right)}\frac{k_{s}}{k_{s}+k_{f}}$ is the signaling-associated parameter group. $\omega_{p}=\beta\mu V_{d}/V_{pms}$is the core-actin rate change that represents the changing rate of core-actin polymerization $\beta\mu V_{d}$ (due to local depolymerization that releases free actin) relative to its steady-state maximum polymerization $V_{pms}$.

The core-actin exchange rate $\omega_{p}$ controls the connection between individual podosome oscillation and spatial wave formation. When $\omega_{p}\to0$, the solution for Eq. S5 is $\left( 1+i\omega\tau_{m} \right)\left( 1+i\omega\tau_{p} \right)-i\omega\tau_{p}\Gamma=0$ or $q^{2}=-{i\omega}/{D_{a}}$; the former solution $\left( 1+i\omega\tau_{m} \right)\left( 1+i\omega\tau_{p} \right)-i\omega\tau_{p}\Gamma=0$ yields the same eigenvalues shown in Eq.8 in the *Models and Methods* using stability analysis on individual podosome dynamics, while the latter solution $q^{2}=-{i\omega}/{D_{a}}$ gives the classical diffusion wave number. As either one can be the solution of Eq. S5, the individual podosome dynamics and spatial diffusion waves in podosome heights are uncorrelated. This is true as our simulations show no wave behaviors in cluster when $\omega_{p}\to0$. In our simulations with parameter values stated in Table S1, we can approximate the relation as:

$$q^{2}=-\frac{i\omega+\omega_{p}}{D_{a}}\left( 1-\frac{\omega_{p}\left( 1+i\omega\tau_{m} \right)}{\left( 1+i\omega\tau_{m} \right)\left( 1+i\omega\tau_{p} \right)-i\omega\tau_{p}\Gamma} \right)\approx-\frac{i\omega+\omega_{p}}{D_{a}}. \left( S6 \right)$$

The above analysis also yields the relation of wavenumber and frequency as shown in Eq.4 in the main text. It is interesting to note that the chemo-mechanical diffusion waves are heavily damped. By defining the phase angle as $\theta=\pi+{tan}^{-1} (\omega/\omega_{p})$, the wavelength can be written as: $\lambda=\frac{2\pi}{\left| \mathcal{R}_{e}\left( q \right) \right|}=\left( \frac{D_{a}^{2}}{{\omega_{p}}^{2}+\omega^{2}} \right)^{\frac{1}{4}}\frac{2\pi}{\cos\left( \theta/2 \right)}$, while the skin or penetration depth that characterizes the damping effect is $\lambda_{s}=\frac{1}{\left| \mathcal{I}_{m}\left( q \right) \right|}=\left( \frac{D_{a}^{2}}{{\omega_{p}}^{2}+\omega^{2}} \right)^{\frac{1}{4}}\frac{1}{\sin\left( \theta/2 \right)}$.
